# Revised evolutionary relationships within Brachycera and the early origin of *bicoid* in flies

**DOI:** 10.1101/2025.07.22.666104

**Authors:** Peter O. Mulhair, Alessandro Pennati, Carlos Herrera-Ubeda, Peter W.H. Holland

## Abstract

The specification of the anterior-posterior axis in the embryo is a crucial step in early insect development. Despite its importance, the underlying genetic and regulatory architecture controlling this process varies significantly between species. In cyclorrhaphan flies, such as *Drosophila melanogaster*, anterior determination is controlled by the transcription factor *bicoid*, which emerged through duplication of the ancestral Hox3 gene (called *zen* in insects). With new, high-quality genomic data we mine the genomes of 186 dipteran species, and find presence of *bicoid* in non-cyclorrhaphan flies, including in the bee flies (family Bombyliidae) and the stiletto flies (Therevidae). We confirm maternal expression and localisation of the non-cyclorrhaphan *bicoid* mRNA to the anterior region of the unfertilised oocyte in the dark edged bee fly, *Bombylius major*. To determine the timing and origin of *bicoid*, we address uncertainty in the dipteran phylogeny, uncovering a ladder-like topology in the branching orders of the early Brachycera lineages. This new species phylogeny suggests that *bicoid* emerged at the common ancestor of Bombyliidae, Asiloidea, and Eremoneura (collectively Heterodactyla), and was subsequently lost at least 16 times. These findings expand our understanding of the early developmental processes in flies and provide new insights into the backbone phylogeny of Diptera and the evolution of *bicoid*.

## Introduction

The insect body plan, including the anterior (head) and posterior (tail) regions, are specified early on during embryogenesis. This process is determined by a network of transcription factors, which activate a cascade response resulting in temporal and spatial expression followed by the differentiation of cells specific to certain regions of the embryo. The exact genes used to regulate this process can vary greatly across insects, and even between closely related species. In Diptera, the anterior determinant (AD) genes, which establish the embryo’s head-to-tail polarity, differ across the order. In the fruitfly, *Drosophila melanogaster*, anterior-posterior (AP) polarity is initiated by *bicoid* ^1^, a gene which has been studied extensively in developmental biology as encoding a morphogen, and in evolutionary biology as revealing the evolution of a transcription factor gene regulatory network ^2–7^. In non-cyclorrhaphan flies, this process is controlled by other, non-homologous transcription factors, such as *cucoid* in culicine mosquitoes, *panish* in midges, *pangolin* in craneflies, and *odd-paired* in moth flies ^8,9^. While there is clear flexibility in which transcription factors adopt these crucial roles in determining AP polarity across flies, there remains large gaps in the fly phylogeny where the presence or absence of these AD genes has not yet been described.

The *bicoid* gene is the primary AD gene within Cyclorrhapha ^10–12^, a large and diverse clade of flies to which *Drosophila melanogaster* belongs. The gene is maternally expressed in nurse cells which deposit RNA into the oocyte; the *bcd* RNA becomes localised anteriorly in the fly egg through active transport along microtubules and an incompletely understood tethering process ^13^. After translation, the Bcd protein then forms a gradient along the AP axis ^1,14^ regulating transcription of a range of target genes ^15–17^. Embryos which lack *bicoid* result in a failure to develop head and thoracic structures, leading to a double abdomen phenotype ^18^. The *bicoid* gene evolved from a tandem gene duplication event of the Hox3 (*zerknüllt* or *zen*) ortholog in flies ^19^; *zen* is involved in the development of the extraembryonic tissue (amnioserosa in cyclorrhaphan flies) ^20^, a function thought to be ancestral to winged insects ^21^. The ancestral *bicoid* gene gained new developmental functions through changes in its 3’ untranslated sequence resulting in maternal expression and localisation to the anterior of the egg ^22^. In addition to this, Bcd gained key amino acid substitutions within the homeodomain (HD), affecting DNA binding activity and establishing RNA binding abilities, which conferred regulatory control over the transcription factors that specify anterior and posterior structures, in particular repressing translation of RNA encoding the transcription factor Caudal which is involved in determining the posterior end ^23–25^.

Functional assays have revealed the crucial substitutions within the Bcd homeodomain permitting its novel patterning activities following duplication. Ancestral sequence reconstruction of both the ancestral Zen homeodomain (AncZB) as well as the ancestral Bicoid homeodomain (AncBcd) of Cyclorrhapha found 31 amino acid differences between the two homeodomains, 11 of which are ‘diagnostic’, present in nine extant Bcd HD sequences and absent from 20 Zen HD sequences ^25^. These ancestral HDs have been tested using rescue assays *in vivo* to assess which changes, or combination of changes, resulted in novel properties. Two key amino acid changes in the recognition helix of the HD, a substitution from glutamine to lysine at position 50 and from methionine to arginine at position 54, when combined, were found to have a large effect in rescuing Bcd’s patterning activity, activating five target genes ^25^. Later work uncovered a complex stepwise route to the evolution of Bcd’s complete patterning activity, which required a combination of substitutions in three subdomains of the HD to ensure robust activity, with the likely existence of suboptimal intermediate Bcd HD sequences during the early evolution of the gene ^7^.

Here, in light of new genomic data across Diptera, we analyse the emergence and evolution of *bicoid*. To accurately determine the timing of origin of the ancestral *bicoid* gene and the subsequent patterns of loss, we require a resolved dipteran species phylogeny. Fly diversity is traditionally separated into two major groups including Nematocera (elongated flies with thin, segmented antennae) and Brachycera (flies with stout bodies and short antennae), with the majority of species within Brachycera belonging to the clade Cyclorrhapha, a group which includes iconic species such as *Drosophila melanogaster*, the house fly, and hoverflies. A robust phylogeny of flies has to this point been difficult to obtain, with limited molecular data resulting in a large amount of uncertainty in the branching patterns between major lineages, particularly in the deeper parts of the phylogeny ^26,27^. The lack of data is confounded by the fact that the early branching lineages within the fly tree represent ancient periods of hyperdiversification, resulting in short branches with low phylogenetic signal ^26^. One of these bursts of diversification occurred at the emergence of Brachycera, which is a large clade consisting of the orthorrhaphous lineages (117 families including more than 100,000 species). A second occurred with the emergence of Cyclorrhapha, a major radiation consisting of more than 64,000 described species from 91 families. The relationships between the non-cyclorrhaphan brachyceran families have undergone multiple variations, most with limited statistical support, including the existence of Orthorrhapha which groups all lineages in a monophyletic group ^26^, and the existence of two monophyletic groups, Homeodactyla (Nemestrinoidea sister to the SXT clade; Stratiomyomorpha, Xylophagidae, and Tabanomorpha) and Heterodactyla (Bombyliidae plus remaining Asiloidea sister to Eremoneura) ^27,28^. Here, we apply our dataset of 186 chromosome level genome assemblies representing 44 families to construct the largest dataset of molecular markers to date used to address the phylogeny of Brachycera dipterans, with a specific focus on the non-cyclorrhaphan families. Our findings recapitulate previously described clades while finding novel relationships for many of the families. We find support for Heterodactyla, with Bombyliidae sister to the remaining lineages (Asiloidea minus Bombyliidae + Eremoneura), and evidence for a ladder-like topology within the remaining non-cyclorrhaphan groups (i.e. paraphyletic Homeodactyla). This new backbone dipteran phylogeny allows us to confidently place the origin of *bicoid* to the common ancestor of Heterodactyla (Bombyliidae sister to Asiloidea + Eremoneura), between 150-200 million years ago, with at least 16 cases of independent loss within this clade. Furthermore, *in situ* hybridisation suggests that the anterior patterning role of *bicoid* dates to this same early date in dipteran evolution.

## Results

### Evolution of *zerknüllt* and *bicoid* across Diptera

We annotated all Hox genes in the genomes of 186 dipteran species (Supplementary table S1), which were represented by chromosome level assemblies thus aiding accurate identification and classification of all genes in the Hox gene cluster ^29^. Focusing on the Hox3 locus (*zen*) and its paralogs (including *bicoid*), we mapped the patterns of gain, loss, and copy number variation across the species tree (Figure 1A). We find extreme variation in the copy number of *zen* across the dipteran phylogeny, with similar rates of duplication found previously in Lepidoptera ^30^, ranging from a single *zen* gene in the crane flies (Tipulidae) to 118 *zen* loci in *Tachina fera* (Tachinidae). We note that this complement of Hox3 loci annotated here may include functional genes, genes containing exon duplications, and pseudogenes. Contrary to previous studies, we find evidence for the presence of *bicoid* in non-cyclorrhaphan lineages, including all four species in the family Bombyliidae (*Anthrax anthrax, Villa cingulata, Bombylius major*, and *Bombylius discolor*) and both species from the family Therevidae (*Thereva unica* and *Thereva nobilitata*) (Figure 1A, Supplementary table S1). The putative *bicoid* sequences from these species group in a clade with the cyclorrhaphan *bicoid* sequences, to the exclusion of all other *zen* sequences, in the gene tree (Figure 1B). There is a distinct long branch separating all *bicoid* sequences from the remaining *zen* sequences in the gene tree, pointing to the high rate of sequence change after the duplication which resulted in *bicoid* (Figure 1B, Supplementary figure S1) ^25^. Interestingly, the length of the branch separating the non-cyclorrhaphan *bicoid* sequences from the remaining cyclorrhaphan branches, albeit not as long as the branch leading to the *bicoid* clade, suggests further accumulation of substitutions in the Bcd homeodomain coinciding with the emergence of Cyclorrhapha (Figure 1B).

**Figure 1.**
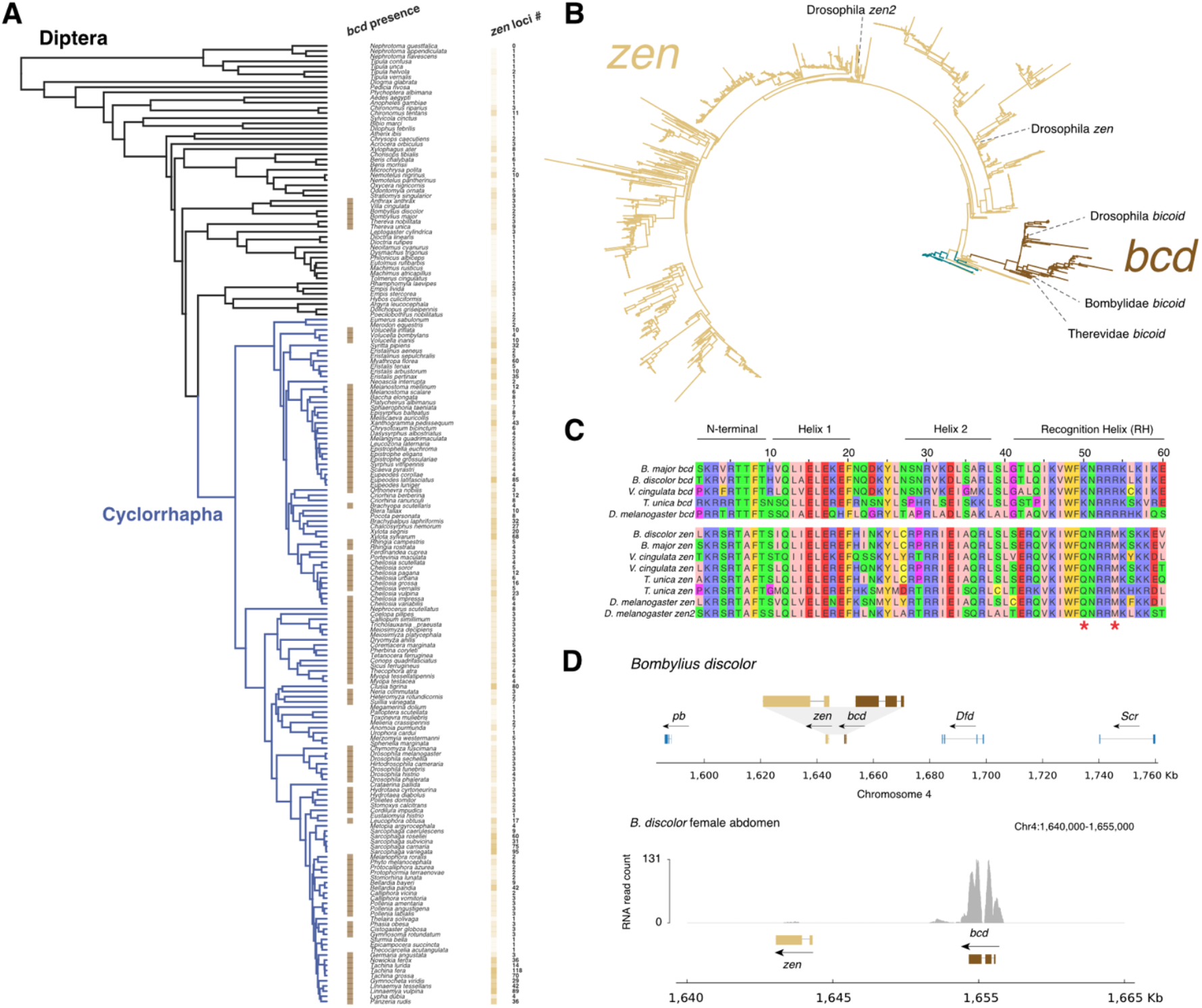
Evolution of zerknüllt (zen) and bicoid (bcd) across the dipteran phylogeny. **(A)** Species tree (left) with the Cyclorrhapha infraorder labelled with blue branches. Presence and copy number (right) of bicoid and zen, respectively. The number of annotated zen loci are represented by colour intensity as well as the number. **(B)** Gene tree of zen (yellow branches) and bcd (brown branches) homeodomain sequences from all dipteran species, including other Hox homeodomain sequences from Apis mellifera, Tribolium castaneum, and Drosophila melanogaster (turquoise branches). **(C)** Multiple sequence alignment of bicoid (upper) and zen (lower) homeodomains from representative non-cyclorrhaphan (Bombylius major, Bombylius discolor, and Thereva unica) and cyclorrhaphan (Drosophila melanogaster) species. Regions of the homeodomain are labelled, including N-terminal, Helix 1, Helix 2, and the Recognition Helix. Red asterisks indicate the key amino acid substitutions at position 50 and 54, shared between all bicoid sequences, which have been previously found to be largely responsible for bicoid novel function in anterior determination. **(D)** Gene tracks (upper) and evidence of expression (lower) for bicoid (brown) and surrounding regions on chromosome 4 of the non-cyclorrhaphan fly Bomylius discolor (Bombyliidae).

Further support for the existence of *bicoid* in Bombyliidae and Therevidae is obtained by examining the alignment of the 60 amino acid homeodomain region with other well-characterised *bicoid* sequences (e.g. from *Drosophila*). The sequences we annotate as *bicoid* in the non-cyclorhaphan lineages share key amino acid substitutions, namely Q50K and M54R (Figure 1C), which are unique to Bcd and have been shown *in vivo* to be crucial for its role in anterior determination ^25^. Finally, we also find that the *bicoid* genes in the non-cyclorrhaphan species have complete, uninterrupted open reading frames (Figure 1D, Supplementary figure S2). We also mapped publicly available RNA sequencing data from the abdomens of a male *Bombylius major* and a female *Bombylius discolor* (Supplementary figure S2) and find expression of *bicoid* to be present only in the *B. discolor* sample (Figure 1D), as we would expect given that *bicoid* is maternally expressed in the abdomen of females where the eggs develop. Taken together, this is strong evidence for the existence of *bicoid* in flies outside of Cyclorrhapha, implying that this gene emerged from a duplication event ∼10-50 million years earlier than previously thought.

### Evidence for anterior localization of *bicoid* mRNA in *Bombylius major* embryo

To determine whether the non-cyclorrhaphan *bicoid* gene possesses maternal expression and anterior RNA localisation, we carried out whole mount *in situ* hybridisation on ovaries dissected from female *Bombylius major* flies (Figure 2). Maternal *bicoid* RNA was detected in unfertilised *B. major* oocytes, tightly localised in an anterior stripe (Figure 2A & B). This pattern of RNA localisation closely mirrors that of *Drosophila melanogaster*, where RNA is transported from nurse cells and becomes tethered at the anterior of the oocyte ^13^. The similar anterior RNA localisation in *Drosophila* and *Bombylius* implies that our identification of *bcd* based on amino acid sequence characters, molecular phylogenetics, and chromosomal location is likely also mirrored in function. We infer that Bcd evolved as an anterior determination in Diptera much earlier than previously reported, before the divergence of Bombyliidae and Cyclorrhapha.

**Figure 2.**
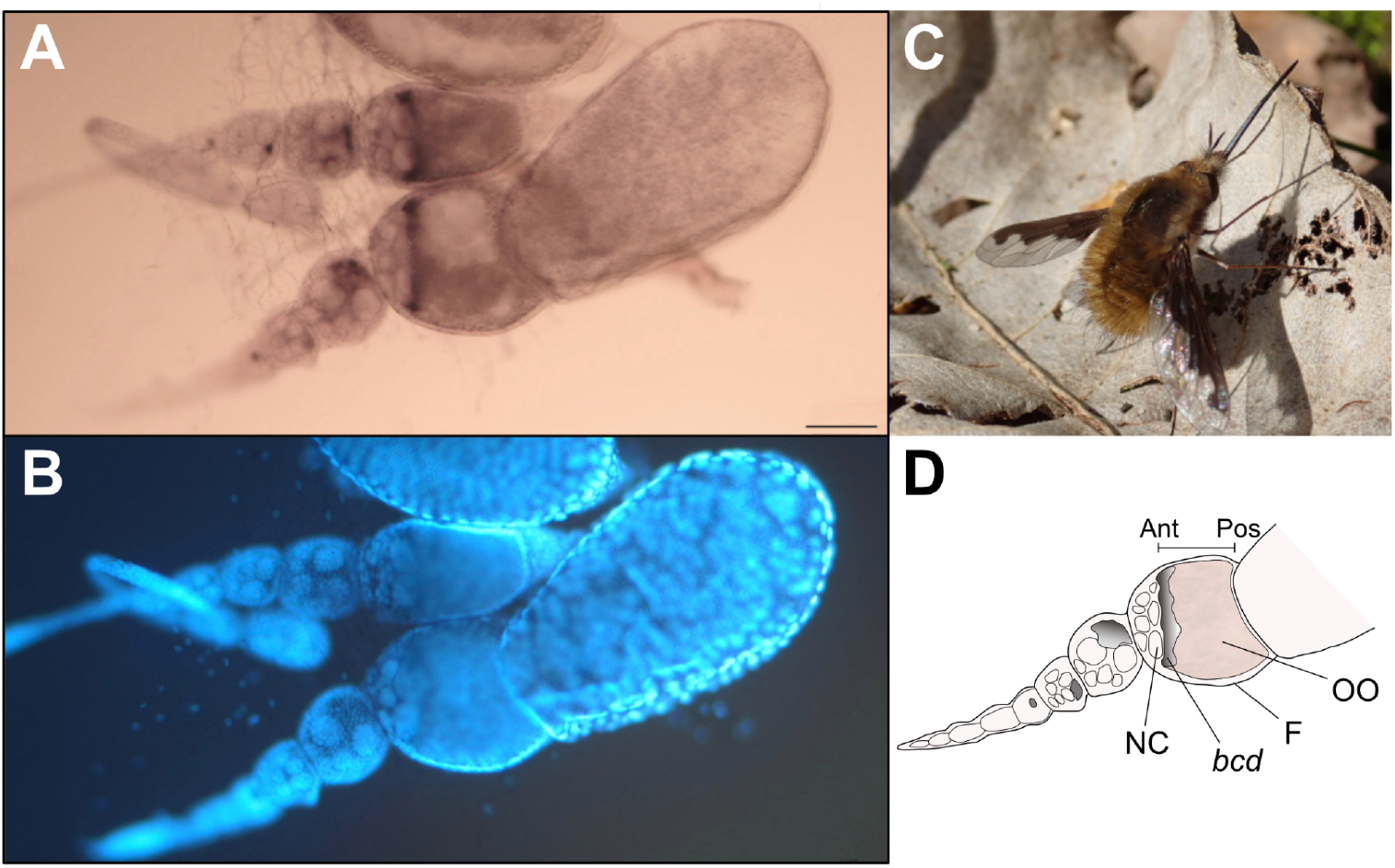
Expression profile of B. major bicoid. **(A)** In situ hybridisation showing RNA distribution of B. major bcd in ovarioles. **(B)** DAPI stain of the same field of view as in (A). **(C)** Female Bombylius major photo taken in the field showing the easily identified female-specific gap between the compound eyes. Photo provided by Dr Liam Crowley. **(D)** Diagram of individual ovariole taken from **(A)** with the AP axis of the oocyte shown and main components labelled; NC=Nurse cells, bcd=bicoid RNA, F=Follicle cells, OO=Oocyte. Scale bar, 100 μm.

### Revisiting deep branching patterns in the Diptera phylogeny

To reconstruct the history of emergence and loss of *bicoid* across the dipteran phylogeny, we first had to address outstanding uncertainty within the brachyceran phylogeny. The dipteran species tree has undergone significant changes with varying degrees of support for clades, particularly within the non-cyclorrhaphan brachyceran lineages (Figure 3A). These brachyceran lineages are composed of 24 described families and include groups such as the soldier flies (Stratiomyidae), robber flies (Asilidae), and bee flies (Bombyliidae). Resolution has been difficult to achieve mainly due to the absence of high quality genomic data across these families, with most studies relying on a small set of genes sampled across a large number of taxa to address this phylogenetic question ^26,27,31^. This is confounded by rapid diversification in early brachyceran evolution ∼200 million years ago, thought to have co-occurred alongside the evolution of flowering plants ^26^. Previous studies grouped almost all non-cyclorrhaphan Brachycera families in a monophyletic group called Orthorrhapha, although support for this clade from morphological and molecular analyses has always been limited ^27,28,32–34^. Strong support exists for the grouping of Empidoidea (including dances flies) with Cyclorrhapha, known as Eremoneura ^28^ (Figure 3A). For the remaining non-cyclorrhaphan lineages, previous work found weak support for a grouping of the three infraorders Stratiomyomorpha (families Stratiomyidae and Xylomyidae), Xylophagomorpha (Xylophagidae), and Tabanomorpha (including Tabanidae and Rhagionidae), known as the SXT clade ^26,27^, and conflicting placement of Bombyliidae, either sister to the remaining Asiloidea families (Therevidea and Asilidae) or as sister to Asiloidea and Eremoneura ^26,27^ (Figure 3A). Here, we revisit the branching patterns of non-cyclorrhaphan brachyceran flies using a taxon sampling which covers most of the families in this part of the tree and a large number of genes. Our dataset consists of species representing the early branching brachyceran families Athericidae (1 species), Tabanidae (1 species), Acroceridae (1 species), Xylophagidae (1 species), Stratiomyidae (9 species), Bombyliidae (4 species), Therevidae (2 species), Asilidae (10 species), Empididae (3 species), Hybotidae (1 species), and Dolichopodidae (3 species), along with 17 outgroup species (representing 8 families) and 133 species from Cyclorrhapha (representing 24 families) (Supplementary table S1). Importantly, these species are represented by chromosome level assemblies, allowing the extraction of a large set of single copy orthologous genes (SCOs) to address this phylogenetic question.

**Figure 3.**
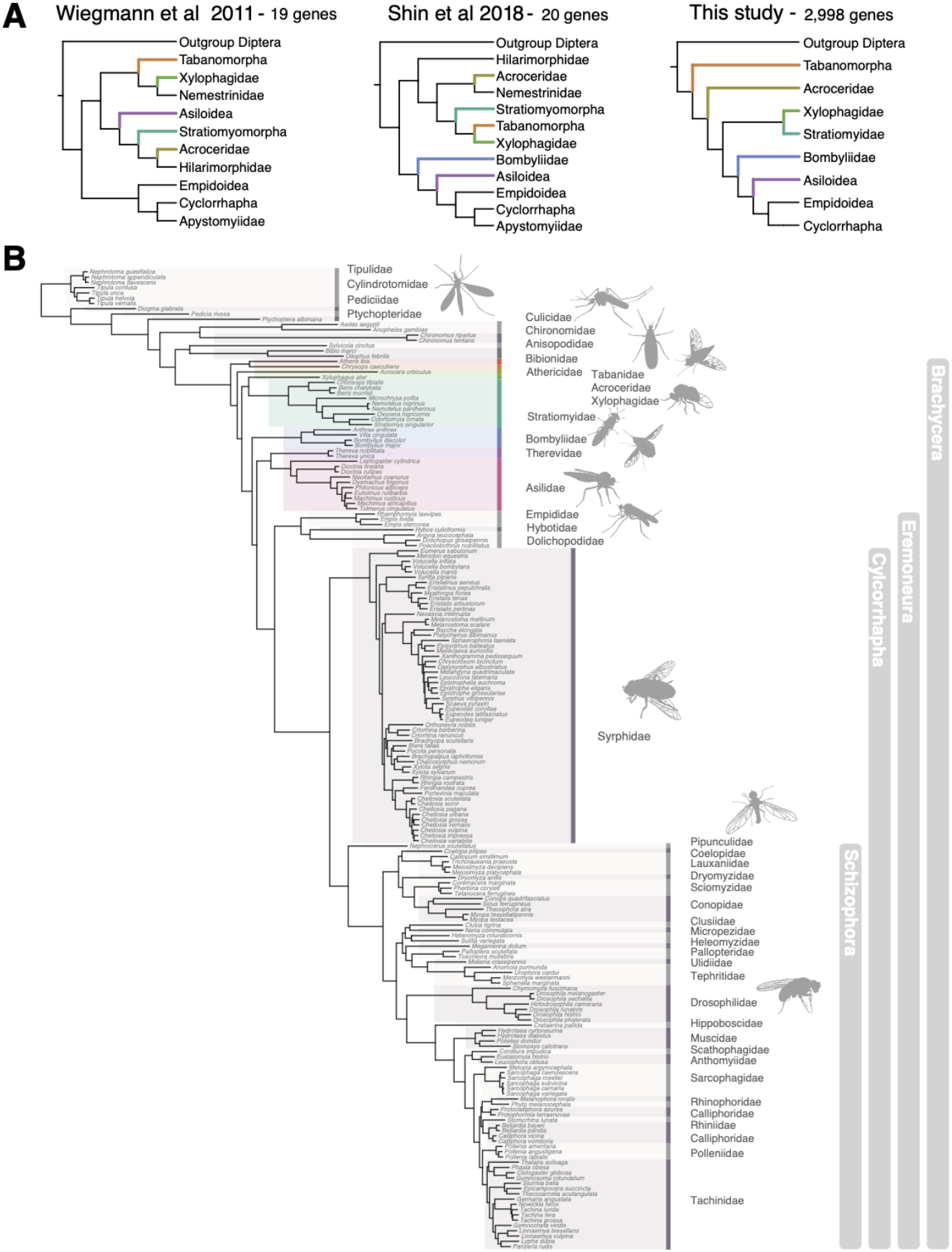
Revised Diptera phylogeny. **(A)** Summary of the high order relationships proposed for the Diptera tree of life from previous studies as well as the updated proposed topology in this study (right). **(B)** Updated species topology for Diptera based on 2,998 genes and inferred under Maximum Likelihood. Families are alternatively coloured and labelled, with specific colours applied to the non-cyclorrhaphan brachyceran lineages that match branch colours in (A). Major groupings within the fly phylogeny are labelled with grey boxes to the right, including Brachycera, Eremoneura, Cyclorrhapha, and Schizophora.

Using the BUSCO (v5.6.1) gene set for Diptera (diptera_odb10) ^35^, we identified 412 single copy orthologs (SCOs) present in all 186 species in our dataset (dat100), and 2,998 SCOs present in at least 90% of the species (dat90). An additional dataset subsetted from the 412 SCOs present in all species was constructed to select genes with greater phylogenetic signal by filtering based on seven alignment and gene tree variables ^36^. Enrichment for genes with these traits has previously been shown to increase phylogenetic signal, particularly for deep branches, and improve overall topological support ^36,37^. Extracting the highest-ranked 70% of such genes resulted in a dataset of 288 SCOs present in all species (datPhy). Finally, we constructed a dataset of BUSCO genes present in all species of a reduced dataset (49 species in total) and enriched for phylogenetic signal, in order to assess the impact of taxon sampling on species tree inference, in particular after removing fast evolving lineages. This last approach resulted in two additional datasets, based on the cutoff of the phylogenetic informativeness test (top 10% ranked genes and top 70% ranked genes as before), of 117 SCOs (datTax10) and 821 SCOs (datTax70) present in all species, with a significantly reduced number of cyclorrhaphan species (9 instead of 133) and a reduced set of non-cyclorrhaphan species (32 instead of 36). In particular, we removed non-brachyceran flies with long branches in the species tree (e.g. *Chironomus* midges) to help reduce the impact of artifacts such as long branch attraction ^38,39^.

The five datasets (dat100, dat90, datPhy, datTax10, datTax70) were then used to infer the species topology applying both concatenation (under Maximum Likelihood) and coalescence based approaches. Across all datasets we recover the monophyly of all dipteran families as expected, a monophyletic Tabanomorpha (Athericidae+Tabanidae, in our dataset), a monophyletic Eremoneura (Empidoidea+Cyclorrhapha) and a monophyletic Cyclorrhapha (Figure 3B, Supplementary figure S3). The coalescence approach recovers a traditional monophyletic Asiloidea (Bombyliidae+Therevidae+Asilidae), however support for this clade reduces when using the datasets consisting of genes with greater phylogenetic signal (datPhy: 0.82, Supplementary figure S3), In addition to this, both datasets with reduced taxon sampling (datTax10 and datTax70) recover full support for the grouping of Bombyliidae sister to Asiloidea (Therevidea and Asilidae) plus Eremoneura (Supplementary figure S4). In all concatenation based trees, we also recover full support for this grouping of Asiloidea excluding Bombyliidae sister to Eremoneura (Figure 3B, Supplementary figure S3 & 4). The grouping of these three clades, known as Heterodactyla, describes brachyceran flies with the presence of a setiform empodium ^34^. This placement of Bombyliidae sister to the remaining clades has been previously recovered, albeit with low support in those studies ^26,27,40^. Within the remaining brachyceran lineages, we consistently recover the SXT clade as paraphyletic, with the Tabanomorpha clade (Athericidae + Tabanidae in our dataset) placed sister to all remaining brachyceran lineages in all analyses (Supplementary figure S3 & 4). Acroceridae (the only family in our dataset representing Nemestrinoidea) is placed as sister to all remaining brachyceran lineages to the exclusion of Tabanomorpha in each of the supermatrix trees, except for datTax10 where it is sister to Xylophagidae, with this grouping branching next to Bombyliidae, Asiloidea, and Eremoneura (Supplementary figure S4). In all but one of the coalescence trees Acroceridae is placed sister to Xylophagidae+Stratiomyidae, while the datTax10 coalescence tree recovers Acroceridae on its own sister to Bombyliidae, Asiloidea, and Eremoneura (classically named Muscomorpha). However, many of the coalescence trees recover these branches with low support (datPhy; 0.75, datTax70; 0.75, datTax10; 0.55). Finally, in all trees except for datTax10 concatenation tree, we recover Xylophagidae as sister to Stratiomyidae, with this clade (including or excluding Acroceridae) placed sister to Heterodactyla (Bombyliidae, Asiloidea, and Eremoneura). Our final tree with the most consistent support thus represents a ladder-like topology within the non-cyclorrhaphan brachyceran lineages, particularly outside of the Heterodactyla clade (Figure 3). Further detailed work is required to resolve the early branching patterns within Brachycera, which will require the inclusion of key lineages absent from this study (Nemestrinidae, Xylomyidae, Hilarimorphidae) as well as more data for undersampled lineages (Athericidae, Tabanidae, and Xylophagidae). This will help to confidently resolve the placement of Nemestrinoidea and the families within the previously hypothesised SXT clade, in particular. Importantly for the purposes of this study, we have confidently resolved the topology of the clades containing the *bicoid* gene (i.e. Bombyliidae, Therevidae, and Eremoneura), allowing for accurate interpretation of the emergence and evolution of the gene.

### Origin, sequence evolution, and loss of *bicoid* across Brachycera

Our findings imply that the ancestor of the brachyceran Zen and Bicoid proteins (AncZB) duplicated, leading to the emergence of ancestral *bicoid* (AncBcd), in the common ancestor of Bombyliidae, Asiloidea, and Eremoneura. This pushes back the time of origin of *bicoid* to between 150-200 million years ago; hence, a large extent of the evolutionary history of *bicoid* has been previously undescribed. This has important implications to understanding of the role of specific amino acid substitutions within the AncBcd homeodomain (HD) which resulted in Bcd’s novel role as a key regulator of anterior determination. To uncover the patterns of substitutions within the AncBcd HD, particularly in the period of time between its origins and the emergence of Cyclorrhapha, we reconstructed ancestral sequences for AncZB HD (common ancestor of all Zen and Bcd sequences) and AncBcd HD (common ancestor of all Bcd sequences, including non-cyclorrhaphan lineages) using a set of insect and fly sequences applied previously ^25^ plus the addition of the new non-cyclorrhaphan Bicoid and Zen HD sequences (Figure 4A). Previous work constructing ancestral HD sequences within Cyclorrhapha found that AncZB (hereafter named AncZB-2018) and AncBcd (hereafter named AncBcd-2018) HDs differed at 31 amino acid positions ^25^. Of these, 11 were thought to be ‘diagnostic’, sites conserved in all current Bcd HD sequences and absent in all insect Zen HD sequences analysed. Our updated ancestral sequences reveal 12 amino acid differences between AncZB-2018 and AncZB and 14 amino acid differences between AncBcd-2018 and AncBcd (Supplementary figure S5), highlighting the importance of including these new non-cylcorrhaphan lineages in the ancestral sequence reconstruction step. In total, we find 21 amino acid positions that differ between AncZB and AncBcd, and of these only 2 are conserved in all fly Bcd HD sequences and absent from all Zen HD sequences, i.e. ‘diagnostic’ (Figure 4B & C, Supplementary figure S6); these are the previously described large-effect substitutions, Q50K and M54R ^25^. Of the 31 sites previously found to differ between AncZB-2018 and AncBcd-2018, only 8 remain different between the updated AncZB and AncBcd, implying that the new AncBcd represents an intermediate state between AncZB and AncBcd-2018 (note that AncBcd-2018 is essentially cyclorrhaphan AncBcd). This is reflected in the fact that we find 7 amino acid differences between AncBcd and the cyclorrhaphan ancestral Bcd HD sequence. We also note a long branch leading to and within the syrphid (hoverflies) Bcd homeodomain sequences suggesting further accumulation of substitutions within Cyclorrhapha (Figure 4B).

**Figure 4.**
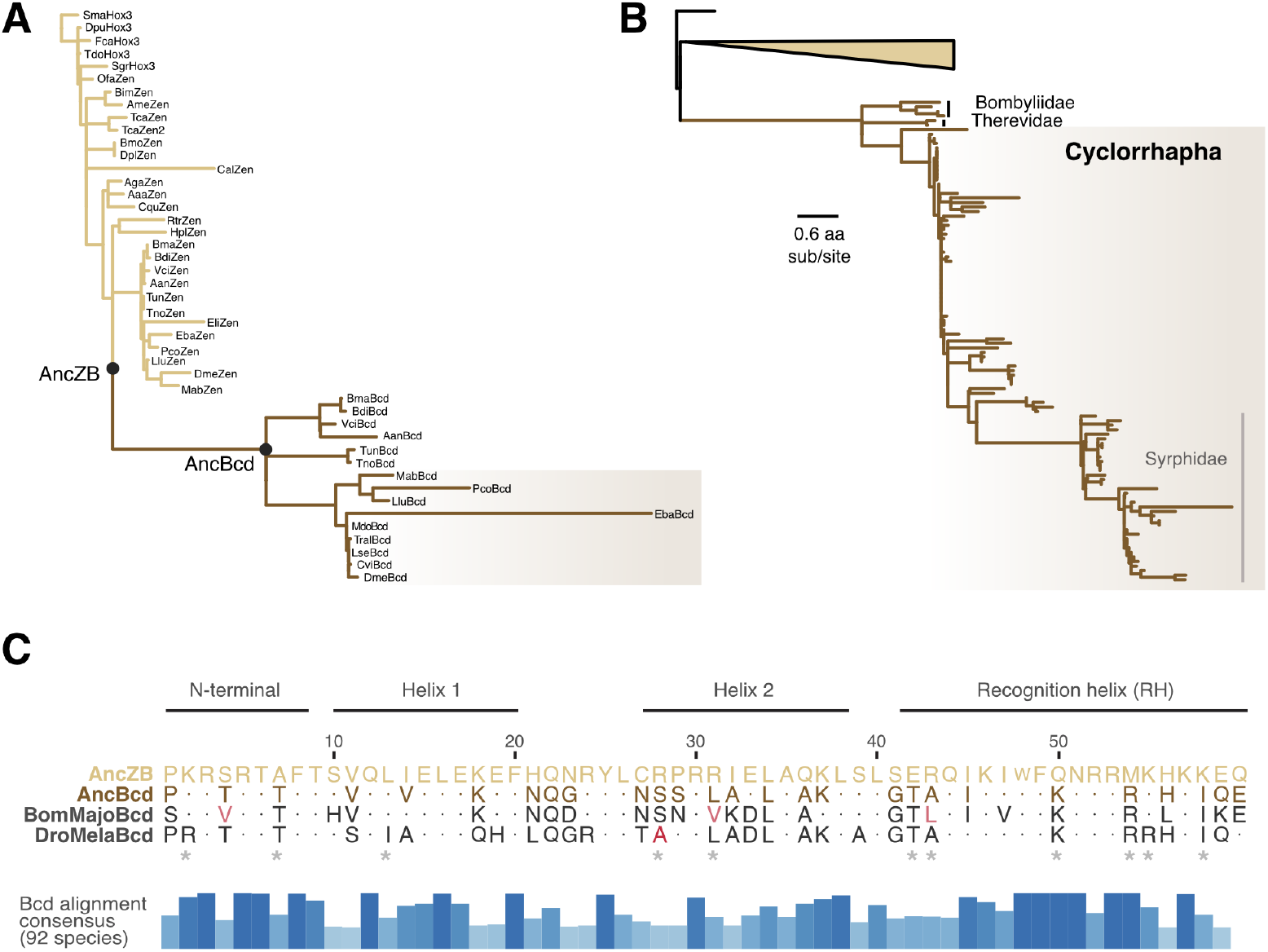
Ancestral sequence reconstruction of Bcd. **(A)** Gene tree of Zen (yellow branches) and Bcd (brown branches) sequences from representative insect lineage (from ^25^) and dipteran species used for ancestral sequence reconstruction. Black nodes represent the reconstructed nodes for the ancestral Bcd homeodomains (AncBcd-HD) and ancestor of all Zen and Bcd homeodomains (AncZB-HD). The Cyclorrhapha clade is labelled with a brown box. **(B)** Expanded gene tree of Zen sequences and Bcd HD sequences from all dipteran species in our dataset. Cyclorrhapha is labelled with a brown box and the families Bombyliidae, Therevidae, and Syrphidae are labelled with text. **(C)** Alignment of the reconstructed amino acid sequences of AncZB and AncBcd, alongside Bombulius major (BomMajo) and Drosophila melanogaster (DroMela) Bcd HD sequences. AncZB is shown as the complete sequence while for AncBcd, BomMajoBcd, and DroMelaBcd, sites that differ from AncZB are shown and conserved sites are shown as dots. Asterisks show the previously designated ‘diagnostic’ sites. The alignment consensus per site across all Bcd sequences in our dataset (92 species) is shown below as blue bar plots, with higher and darker bars representing higher conservation (see Supplementary figure S6 for the full Bcd alignment).

Outside of the HD, we find an even greater extent of sequence and structural divergence in Bicoid. When we align the complete protein sequences of Bicoid for select non-cyclorrhaphan and cyclorrhaphan species, we observe low overall sequence identity between the two groups with higher rates of sequence conservation within the phylogenetic groups (i.e. non-cyclorrhaphan and cyclorrhaphan lineages) (Supplementary figure S7). This implies that while *bicoid* is fast evolving across longer evolutionary distances, there is sequence conservation between closely related species. These divergent rates of evolution within *bicoid* across flies is reflected in the protein structure, where structural alignment between *Bombylius major* and *Drosophila melanogaster* shows strong overlap in the conserved homeodomain region but low structural alignment outside of this, likely due to significant structural disorder in these regions of the protein (Supplementary figure S7).

Our finding that *bicoid* RNA is localised anteriorly in the bee-fly *Bombylius* is particularly intriguing when combined with the deduced species phylogeny and genome analyses identifying subsequent *bicoid* gene losses. The implication is that even after *bicoid* had evolved its role in patterning of the anterior-posterior axis, there were multiple losses of the *bcd* gene. Hence, the developmental pathway for anterior specification is even more evolutionarily labile than previously thought ^8^ (Figure 5, Supplementary figure S8). For example, in the non-cyclorrhaphan brachyceran lineages, we find no evidence for the presence of *bicoid* in the family Asilidae or the infraorder Empidoidea (families Empididae, Hybotidae, and Dolichopodidae) suggesting at least two independent losses of *bicoid* in these lineages (Figure 5). Within Cyclorrhapha, *bicoid* has been lost from multiple families, including several times within Syrphidae (18 out of 56 species missing *bicoid*), Clusiidae (represented by *Clusia tigrina*), Megamerinidae (*Megamerina dolium*), Pallopteridae (missing in both represented species, *Palloptera scutellata* and *Toxonevra muliebris*), Ulidiidae (*Melieria crassipennis*), Tephritidae (missing in all 4 species), Hippoboscidae (*Crataerina pallida*), Anthomyiidae (missing in one of two species), Sarcophagidae (all 6 species), and Tachinidae (4 out of 17 species) (Figure 5). In total, given our current dataset, we estimate at least 16 independent cases of *bicoid* loss throughout Diptera (Supplementary figure S8). All these *bicoid* gene losses occurred after *bicoid* evolved anterior RNA localisation in oocytes.

**Figure 5.**
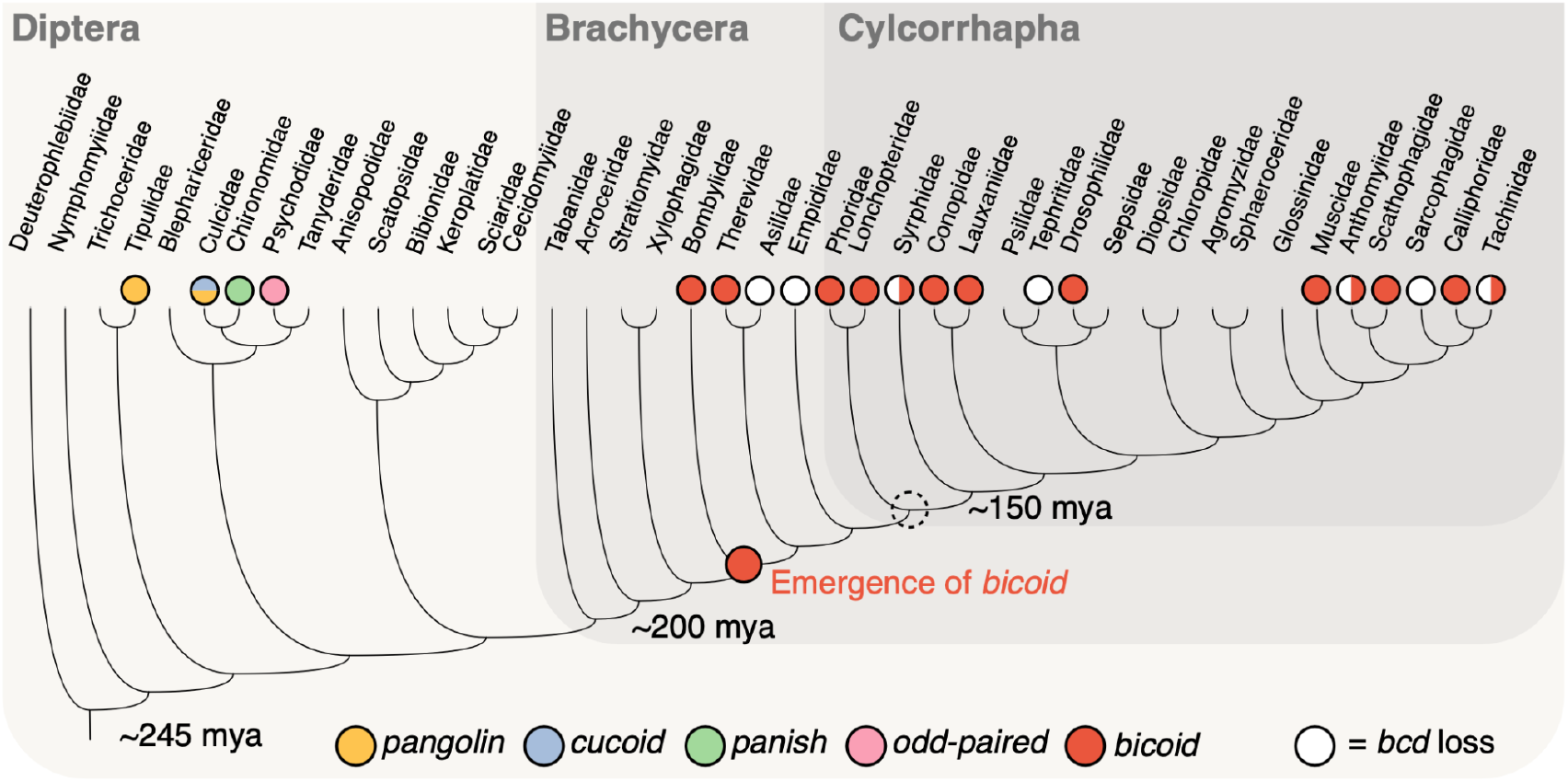
Summary of bicoid evolution across Diptera. The cladogram contains higher level relationships of Diptera families. Select non-brachyceran families are labelled by the presence of previously discovered anterior determinant genes (eg. pangolin in Tipulidae). Bicoid presence and absence is labelled by red and white circles, respectively, on the tips of the tree. Lineages where the presence of bicoid remains unknown have no circle present. The emergence of bicoid is labelled at the common ancestor of Bombyliidae and Asiloidea + Eremoneura. Previously thought node of origin of bicoid, Cyclorrhapha, is labelled with a broken, empty circle.

## Discussion

This study represents the most comprehensive assessment of the evolution of Hox3 genes in flies to date. Annotation of all Hox genes in 186 chromosome-level dipteran genomes from 44 families uncovers dynamic rates of copy number variation in the Hox3 loci, as previously discovered in Lepidoptera and some other insect orders ^30,41^. Our analysis using an expanded sampling of dipteran taxa has, for the first time, revealed the presence of *bicoid* in non-cyclorrhaphan flies, which evolved by tandem duplication from *zen* followed by extensive sequence divergence. We confirm that this newly discovered *bicoid* is homologous to the well described fruit fly *bicoid*, including sharing key functional substitutions which permitted its novel activity (Q50K and M54R). Furthermore, we show that *bicoid* RNA is maternally expressed and localised to the anterior pole of unfertilized oocytes in a non-cyclorrhaphan fly, *Bombylius major* (Bombyliidae). This finding has important implications for our understanding of when *bicoid* emerged, how it gained novel functionality in anterior specification, and the lability of anterior patterning mechanisms in insects.

Previously, it was thought that *bicoid* emerged by duplication of the ancestral *zen* gene at the base of Cyclorrhapha. Detailed molecular work used a reconstructed sequence of the ancestral Bicoid homeodomain ^25^ to generate chimeric homeodomains containing different numbers of ancestral substitutions in the homeodomain. These were then used to carry out gene rescue assays in *Drosophila melanogaster* to determine the order of amino acid substitutions which lead to *bicoid*’s novel function ^7^. This study revealed that combinations of substitutions in at least three subdomains of the homeodomain, not just the two substitutions of large effect in the recognition helix (Q50K and M54R), were required to rescue *bicoid*’s full patterning activity. However, suboptimal patterning activity was achieved with combinations of substitutions in two of the three subdomains suggesting a stepwise route to the robust activity of *bicoid* in extant species ^7^. Our analyses push the origin of *bicoid* deeper in time and give insight into earlier stages in its evolution. Our newly constructed AncBcd HD, deduced using data from non-cyclorrhaphan flies and a much larger sample of dipteran genomes, reveals there are fewer amino acid differences between AncZB (the ancestor of *zen* plus *bicoid*) and the first bicoid (AncBcd), as well as far fewer conserved ‘diagnostic’ sites - those conserved across all extant Bcd HD sequences but absent from Zen HD sequences. In fact, we find just two diagnostic sites, the previously described large effect lysine at position 50 and arginine at position 54 ^25^. This suggests that outside of these two key sites, there is a large amount of plasticity in the core sites involved in *bicoid*’s full patterning activity. The possible combination of key amino acids in different subdomains is now greatly increased, pointing to flexibility in sites determining *bicoid*’s activity across flies.

Not only is there flexibility in the specific amino acids used for anterior embryo patterning, there is clear flexibility in the gene employed to do this across flies ^8,9^. This has been previously described by assessing putative AD genes in a range of non-cyclorrhaphan flies which do not possess *bicoid*, uncovering unrelated genes encoding AD in different fly families (Figure 5), as well as a role for alternative splicing in this process ^9^. Our findings suggest that even after the establishment of *bicoid* as the AD gene, this important developmental step has undergone major shifts in its underlying gene regulatory networks, given that *bicoid* has been lost many times throughout flies (Figure 5). We document at least 16 cases of *bicoid* loss in our dataset, including loss from complete families (e.g. Asilidae, Empididae, and Tephritidae). These losses occurred after *bicoid* was recruited for anterior specification, as inferred by the RNA localisation in *Bombylius*. Further work to fill the gaps in our understanding of the AD genes in these and non-cyclorhaphan families without *bicoid* will be of great interest to understand the true rate of molecular change in this developmental process.

The dipteran phylogeny has long been difficult to resolve in many key nodes, particularly along the backbone of the Brachycera clade. Here we have used a large number of molecular markers, and many more genomes, to resolve some but not all of the controversial phylogenetic nodes. In particular, we focused attention on the part of the phylogeny required to resolve the origin and losses of *bicoid*, specifically between the superfamilies Asiloidea and Empidoidea, as well as their relationship to the Cyclorrhapha. Both concatenation and coalescence based approaches find strong and consistent support for the existence of Heterodactyla (Asiloidea sister to Eremoneura) ^28^, with the family Bombyliidae placed outside of Asiloidea (Asilidae+Therevidae in our dataset) and sister to Asiloidea and Eremoneura (found in all concatenation trees and both taxon reduced coalescence trees with full support; Figure 3). This topology has been recovered before, but never with high support values ^26,27,40^. Interestingly, all coalescence phylogenies, except for the trees with reduced number of species, recover a monophyletic Asiloidea (including Bombyliidae) in all three datasets, but with lower support values, particularly in the dataset enriched for greater phylogenetic signal. This discordance between concatenation and coalescence approaches can result from a range of biological and technical issues such as difficulties in constructing accurate gene trees for deep divergences ^42^, effects from patterns of reticulate evolution such as incomplete lineage sorting, horizontal transfer, or introgression ^43,44^, or simply due to incorrect inference of the species tree from concatenation approaches ^45,46^. The discordance between trees constructed using a reduced number of species (49 species) and the full set of species (186 species) also highlights the impact of taxon sampling in resolving the early branching patterns of Brachycera, in particular the potential impact of including fast evolving outgroup lineages and resulting long branch attraction ^38,39^.

Regardless of the remaining conflict between approaches, the current phylogenetic resolution allows us to confidently place the origin of *bicoid* at the base of Heterodactyla. The remaining non-cyclorrhaphan families, however, proved more difficult to confidently resolve. In all analyses the Homeodactyla clade (SXT clade + Nemestrinoidea) is recovered as non-monophyletic, with the SXT clade itself also never being recovered, a finding which has been recorded previously ^47–49^. Tabanomorpha (represented by Athericidae+Tabanidae in this study) is consistently monophyletic and placed sister to all remaining brachyceran lineages in our dataset with full support. The relationships between Stratiomyidae, Xylophagidae, and Acroceridae remain conflicted between concatenation and coalescence phylogenies, with different topologies often recovered with low support values, in particular in datasets with fewer characters (dat100, datPhy, datTax10, datTax70). Resolving branching patterns of such ancient radiations remains a consistent issue throughout phylogenomic studies, particularly within the diverse holometabolous insect orders ^50^. This difficulty in understanding the true relationships derives from a combination of scarce phylogenetic information as a result of rapid species radiation and the difficulty in extracting the true phylogenetic signal from misleading signal as a result of substitution saturation, or incomplete lineage sorting or hybridisation ^39,51^. Ongoing research is continuing to shed more light on this particular difficult to resolve insect phylogeny with important implications for the taxonomy of the order ^52–55^. More high-quality genomes will provide a greater number and more diverse type of characters to help in its resolution^39–56^.

With a more phylogenetically diverse set of genomes there is ample opportunity, as shown here, to delve deeper into the genetic basis of fly evolution and development. Considering the vast diversity of living flies, there still remain gaps in our understanding of how developmental processes have evolved, and which pathways are more or less prone to evolutionary change. The increased number of dipteran genomes will allow further detailed characterisation of the molecular basis of development and adaptation in this fast evolving and diverse clade of insects.

## Data and code availability

Data and code required to reproduce all results can be found in the Supplemental material and on Figshare (https://figshare.com/s/c47ea5b5716fa439b3ca).

## Acknowledgements

We thank Asia Hoile and Riccardo Kyriacou for help with dissecting bee fly ovaries and Tom Lewin for guidance on experimental design. We are also grateful to all members associated with the Darwin Tree of Life project who carried out all sampling, processing, and sequencing of the samples used in this study. This research was funded by the Wellcome Trust Darwin Tree of Life Discretionary Awards (218328, 226458), the John Fell OUP Research Fund, and the BBSRC Fellowship (grant UKRI893 to P.O.M.).

## Methods

### Genome acquisition and gene annotation

Dipteran genomes were downloaded from NCBI using ncbi datasets ^57^: a list of all genomes analysed are present in Supplementary table S1. All genomes in our dataset are chromosome level, with most produced by the Darwin Tree of Life project; NCBI accession number PRJEB40665, ^58^. The dataset consists of 186 species representing 44 dipteran families. To annotate the Hox genes in all species we used the HbxFinder pipeline (github.com/PeterMulhair/HbxFinder; ^29^). This pipeline uses homeodomain sequences from *D. melanogaster, T. castaneum*, and *Apis mellifera*, obtained from HomeoDB ^59,60^, as queries for iterative sequence similarity searches against genomes. Briefly, a tBLASTn search against the dipteran genomes were carried out using an e-value threshold of 1×10−5 and non-overlapping hits were extracted to retain a single sequence per Hox gene. The resulting dipteran Hox homeodomain sequences were used in a reciprocal BLASTx search against the complete homeodomain protein data set, where hits with significant percentage identity (>70%), provided initial identification of the given Hox gene. Next, another round of sequence similarity searches was performed using MMseqs2 ^61^ and 1kb on either side of the homeobox genes annotated from the initial search. Finally, the newly identified dipteran Hox nucleotide sequences were then translated into an amino acid using the sixpack package from EMBOSS ^62^.

### Gene tree inference and expression quantification

Homeodomain sequences from all Hox genes were used to construct a gene phylogeny. The 60 amino acid sequences were aligned using Mafft v7.467 ^63^ and trimmed using trimAl v1.4.rev15 ^64^ and phylogenetic reconstruction was carried out using IQ-Tree 2.0-rc1 ^65^ applying ModelFinder to find the model of best fit ^66^. Multiple sequence alignments were visualised using Ggmsa ^67^, and gene trees were visualised using Ggtree ^68^. Interrogation of the gene tree allowed for confirmation of the identity of the specific Hox genes. Gene tracks of *zen* and *bicoid* were plotted using pyGenomeTracks ^69^ and gene tracks with expression showing RNA coverage were visualised using trackplot (github.com/PoisonAlien/trackplot; ^70^). RNA-seq data from *Bombylius major* and *Bombylius discolor* were obtained from the Sequence Read Archive on NCBI (ERR10123677 and ERR10123673). RNA from both specimens was obtained from adult abdomens, with sex validated using the photographic evidence of the individual used to generate the sequence data (Supplementary figure S2). RNA reads were trimmed for quality using Trimmomatic v0.39 ^71^, and the processed RNA reads were mapped to each genome using bowtie2 ^72^ in order to confirm evidence of expression and visualise RNA read depth.

### *In situ* hybridisation

The *Bombylius major bicoid* complete open reading frame was synthesised and inserted into pBluescript II KS by Genscript (USA). To generate template for *in vitro* transcription, PCR amplification was carried out using Q5 High-Fidelity DNA Polymerase (NEB, M0491S) and M13/reverse M13 primers followed by purification with the Monarch Spin PCR & DNA Cleanup Kit. (NEB, T1130S). Digoxigenin labelled RNA probe was synthesised using the mMESSAGE mMACHINE T7 Transcription Kit (Invitrogen, AM1344) in combination with DIG RNA Labelling Mix (Roche, 11277073910). Adult female *B. major* were collected by net from Wytham Woods, Oxfordshire, UK (latitude 51.77, longitude –1.34) on 9 April 2023 and killed by freezing. Ovaries were dissected in sterile phosphate buffered saline (PBS) pH 7.4, fixed in 4% paraformaldehyde in PBS and stored in 100% methanol at -20°C for up to 2 years. Whole mount *in situ* hybridization followed the *Drosophila* protocol of ^73^ with slight modifications. Ovaries were rehydrated and treated with 50 µg/ml proteinase K for 1 hour at room temperature, washed with PBS containing 0.1% Tween 20 (PBT) and post-fixed in 4% paraformaldehyde for 1 hour at room temperature. Samples were extensively washed and incubated in hybridization solution (50% formamide, 5× SSC, 50 µg/ml heparin, 100 µg/ml tRNA, 100 µg/ml salmon sperm DNA, 0.1% Tween 20) for 1 hour at 50°C. Hybridization was performed overnight at 50°C in hybridization solution containing 1 µl/ml RNA probe. Samples were washed multiple times in hybridization solution at 65°C, followed by additional washes in PBT, then incubated in PBT containing 1% Blocking Reagent (Roche, 11096176001) and 1:2000 Anti-Digoxigenin-AP (Roche, 11093274910) overnight at 4°C. After further washes in PBT, ovaries were stained using 10 µl/ml NBT/BCIP solution (Roche, 11681451001) in NTMT buffer (0.1 M NaCl, 0.1 M Tris-HCl pH 9.5, 0.05 M MgCl2, and 0.1% Tween 20) and post-fixed in 4% paraformaldehyde. Samples were mounted in VECTASHIELD Antifade Mounting Medium with DAPI (Vector Laboratories, H-1200) and imaged using a Zeiss Axioscope 2 Plus microscope equipped with a ChromyxHD camera.

### Phylogenetic analysis

Phylogenetic reconstruction was carried out on conserved genes annotated by BUSCO v5.3.2 ^35^ using the Diptera BUSCO gene set (diptera_odb10). BUSCO completeness scores were generally high across all species (ranging from 88.9%-99.3%, mean 96.7% BUSCO completeness), reflecting the high quality of the underlying genomes used in this analysis. The resulting data consisted of 412 genes present in single copy across all species (dat100) and 2,998 genes present in at least 90% of the species (dat90). These datasets were filtered to enrich for genes with greater phylogenetic signal using genesortR ^36^. In addition to this, a dataset consisting of a reduced taxon dataset, in particular removing fast evolving non-brachyceran outgroup lineages and cyclorrhaphan lineages, was produced to assess the impact of taxon sampling on tree inference. For each gene in all datasets, sequences were aligned using Mafft and trimmed using ClipKIT v1.3 ^74^. Individual gene trees were constructed using IQ-Tree2 with the best fit substitution model automatically selected by ModelFinder and with 1,000 bootstrap replicates. Concatenated alignments were constructed from all datasets (dat100, dat90, datPhy, datTax10, datTax70) using PhyKIT ^75^. Species trees were inferred from these data using IQ-Tree2 for maximum likelihood based inference applying the LG matrix and a discrete gamma rate model with four categories (LG+G4+I). Coalescence-based species trees were constructed with ASTRAL v5.7.7 ^76,77^ using gene trees inferred from the trimmed BUSCO gene alignments as described above. Species trees were plotted using Toytree ^78^.

### Ancestral sequence reconstruction

In order to compare our results to previous findings, ancestral sequence reconstruction was carried out using Zen and Bcd HD sequences from a set of 27 hexapod species previously used ^25^ along with the additional non-cyclorrhaphan sequences. Amino acid sequences were aligned using MAFFT and a constrained gene tree was constructed using IQ-Tree2, with a topology reflecting correct species relationships. Ancestral sequence reconstruction of the homeodomains was carried out using a maximum likelihood method implemented in the codeml package of PAML v4.10.6 ^79^. This was carried out with the constrained gene tree, using the LG model, and four-category gamma distribution of among-site rate variation. Per site confidence was assessed by measuring the posterior probabilities; amino acid sites where multiple states have a PP > 0.2 were labelled as low confidence. The ancestral Zen-Bcd HD (AncZB) and ancestral Bcd HD (AncBcd) were aligned to modern Zen and Bcd HD sequences from all species in our dataset to determine diagnostic sites; sites that differ between AncZB and AncBcd and are present in all modern Bcd HD sequences and absent from Zen HD sequences. The full open reading frames for *Drosophila melanogaster* and *Bombylius major* Bicoid were extracted and protein structural models were built using Alphafold ^80^ implemented in the command line tool gget ^81^.

