## Supplemental_Material for "Revised evolutionary relationships within Brachycera and the early origin of *bicoid* in flies"

#### Supplementary Figures

Supplementary Figure S1: Gene tree of all *zen* and *bcd* homeodomain sequences.
Supplementary Figure S2: Gene tracks and expression of *zen* and *bcd* genes in Bombyliidae.
Supplementary Figure S3: Summary of phylogenetic trees constructed using different datasets and approaches.
Supplementary Figure S4: Summary of phylogenetic trees built using the reduced taxon dataset.
Supplementary Figure S5: Updated ancestral sequence reconstruction of AncZB and AncBcd.
Supplemental Figure S6: Alignment of Zen and Bcd homeodomains across all species.
Supplemental Figure S7: Evolution of complete *bicoid* ORF sequences.
Supplementary Figure S8: Patterns of *bicoid* loss across flies.

#### Supplementary Tables

Supplementary Table S1: Table of genomes used in this study, including accession number and genome citation.

#### Supplementary References

Reference list for genomes listed in Supplementary Table S1.

#### Supplementary Figures

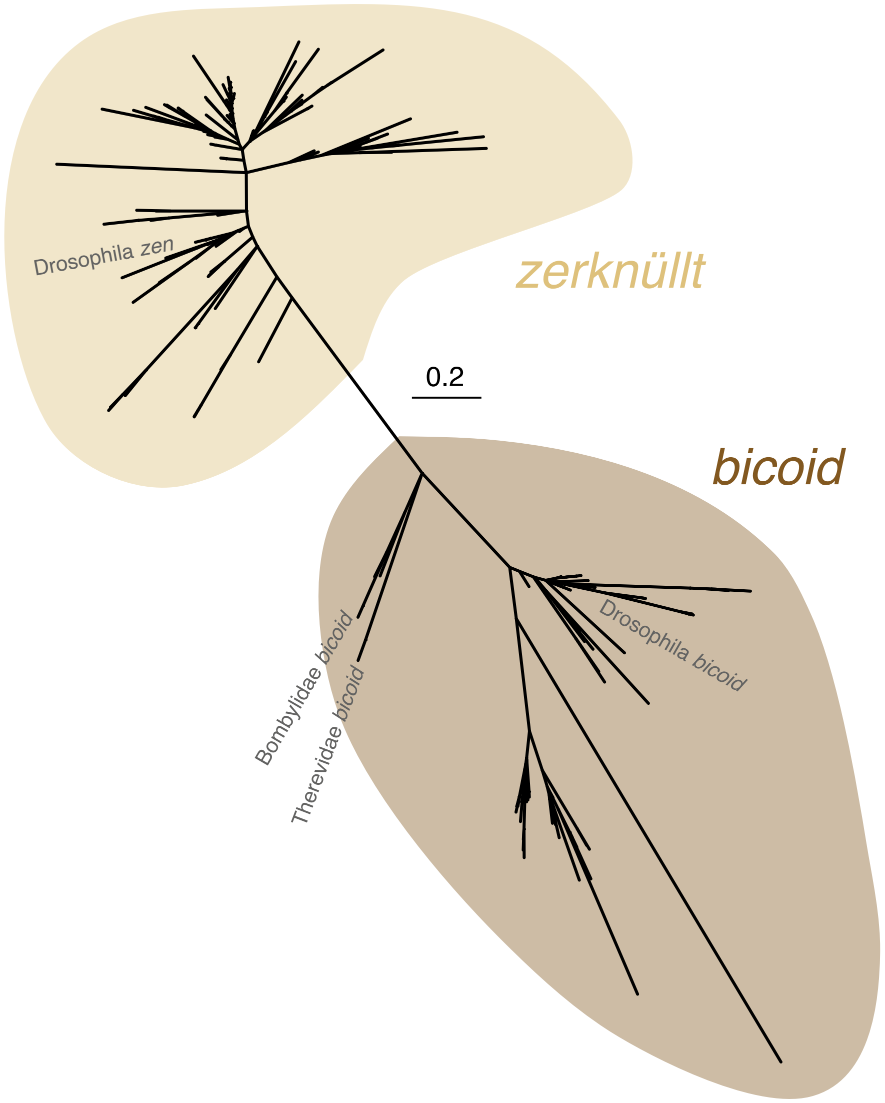


**Supplemental Figure S1: Gene tree of all *zen* and *bcd* homeodomain sequences.** *Zen* branches are labelled and coloured in a yellow colour while *bcd* branches are labelled and coloured in brown. *Drosophila melanogaster zen* and *bicoid*, as well as the non-cyclorrhaphan *bicoid* sequences are labelled. A long branch separates the *zen* and *bcd* clades.

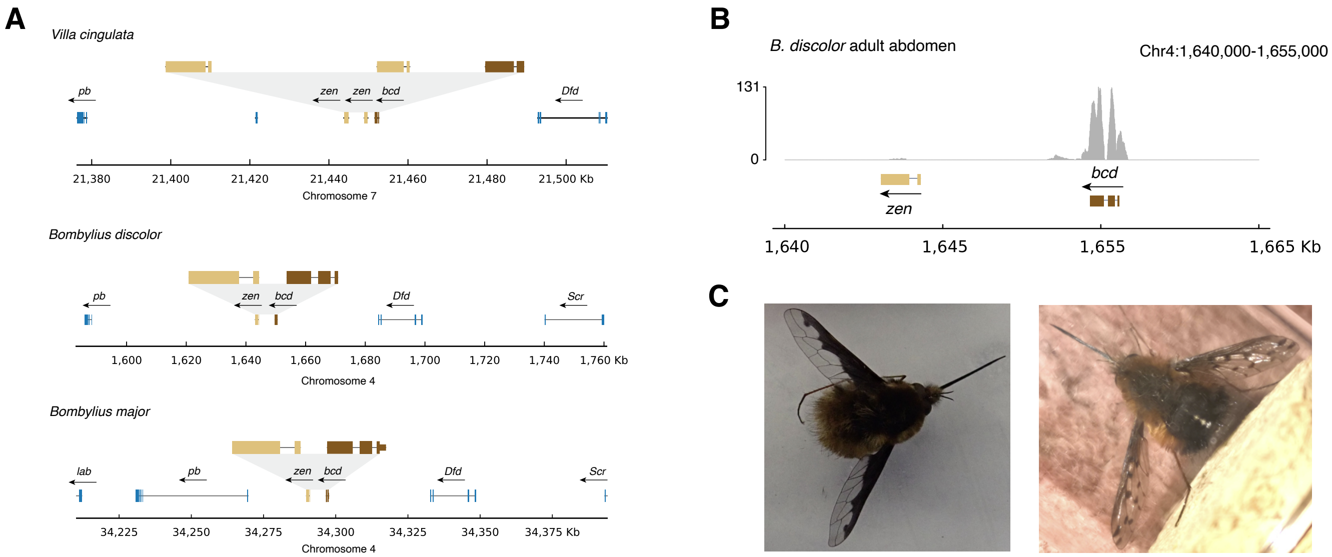


**Supplemental Figure S2: Gene tracks and expression of *zen* and *bcd* genes in Bombyliidae. (A)** Gene models for *zen* and *bcd*, as well as the other Hox genes immediately up- and downstream are shown for three Bombyliidae species for which gene annotation data was available; *Villa cingulata*, *Bombylius discolor*, and *Bombylius major*. The chromosome number and position in each species is given on the x-axis, as well as the orientation and intron/exon structure for each of the genes. **(B)** Evidence of expression of *bcd* in *Bombylius discolor*. Peaks along the y-axis represent the number of RNA reads from *Bombylius discolor* female adult abdomen which map to the *bcd* loci. **(C)** Photographs of the individuals sampled and used for genome and RNA sequencing by the Darwin Tree of Life project for *Bombylius major* and *Bomylius discolor*, respectively.

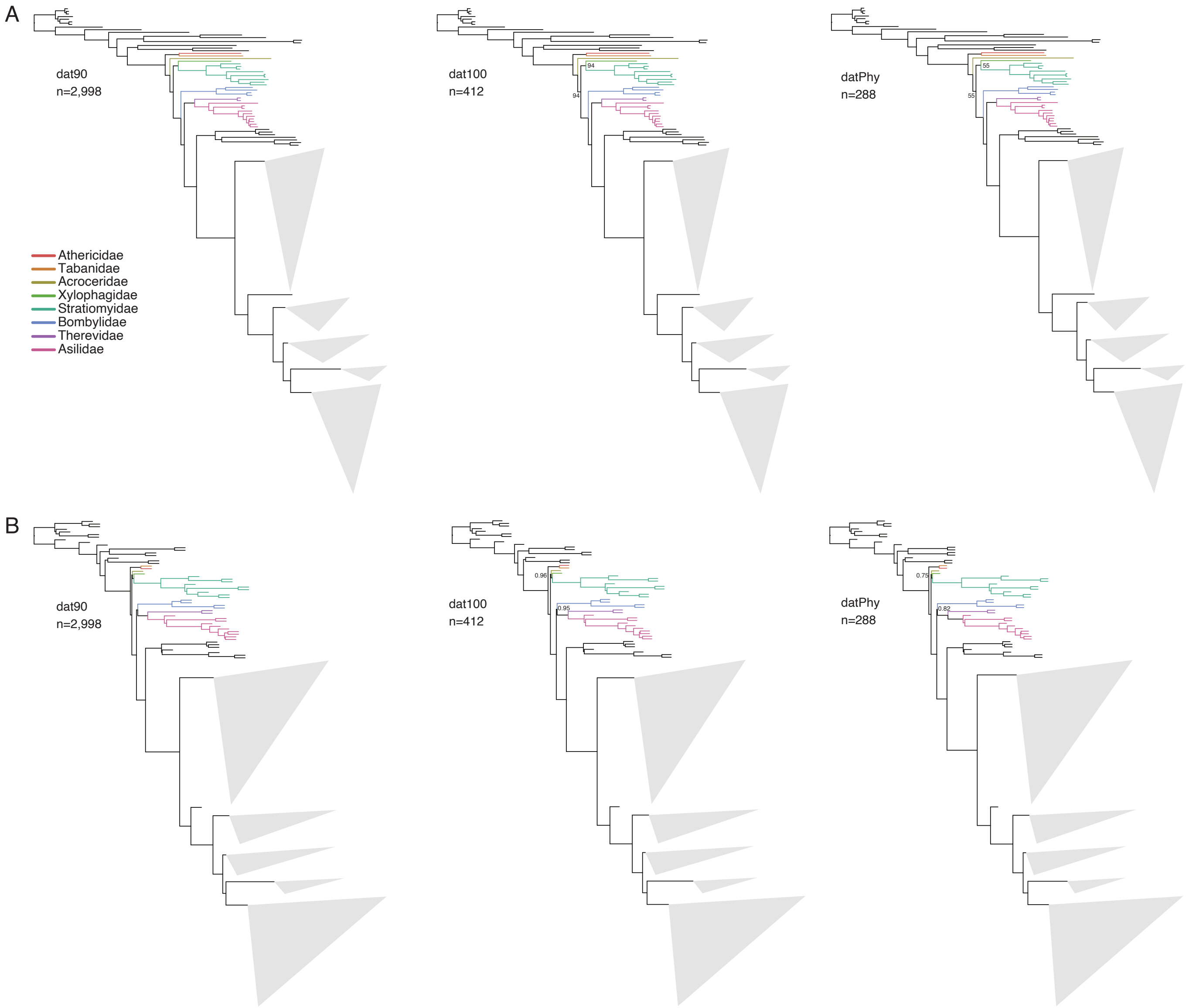


**Supplemental Figure S3: Summary of phylogenetic trees constructed using different datasets and approaches.** In all phylogenies, the non-cyclorrhaphan brachyceran branches are coloured according to the key on the left, as in Figure 3. Major cyclorrhaphan clades are collapsed into grey triangles for visualisation purposes. **(A)** Phylogenetic trees constructed using concatenated matrix of dat90, dat100, and datPhy under a maximum likelihood approach **(B)** Phylogenetic trees from the same datasets as above, constructed instead using gene trees and a coalescence approach.

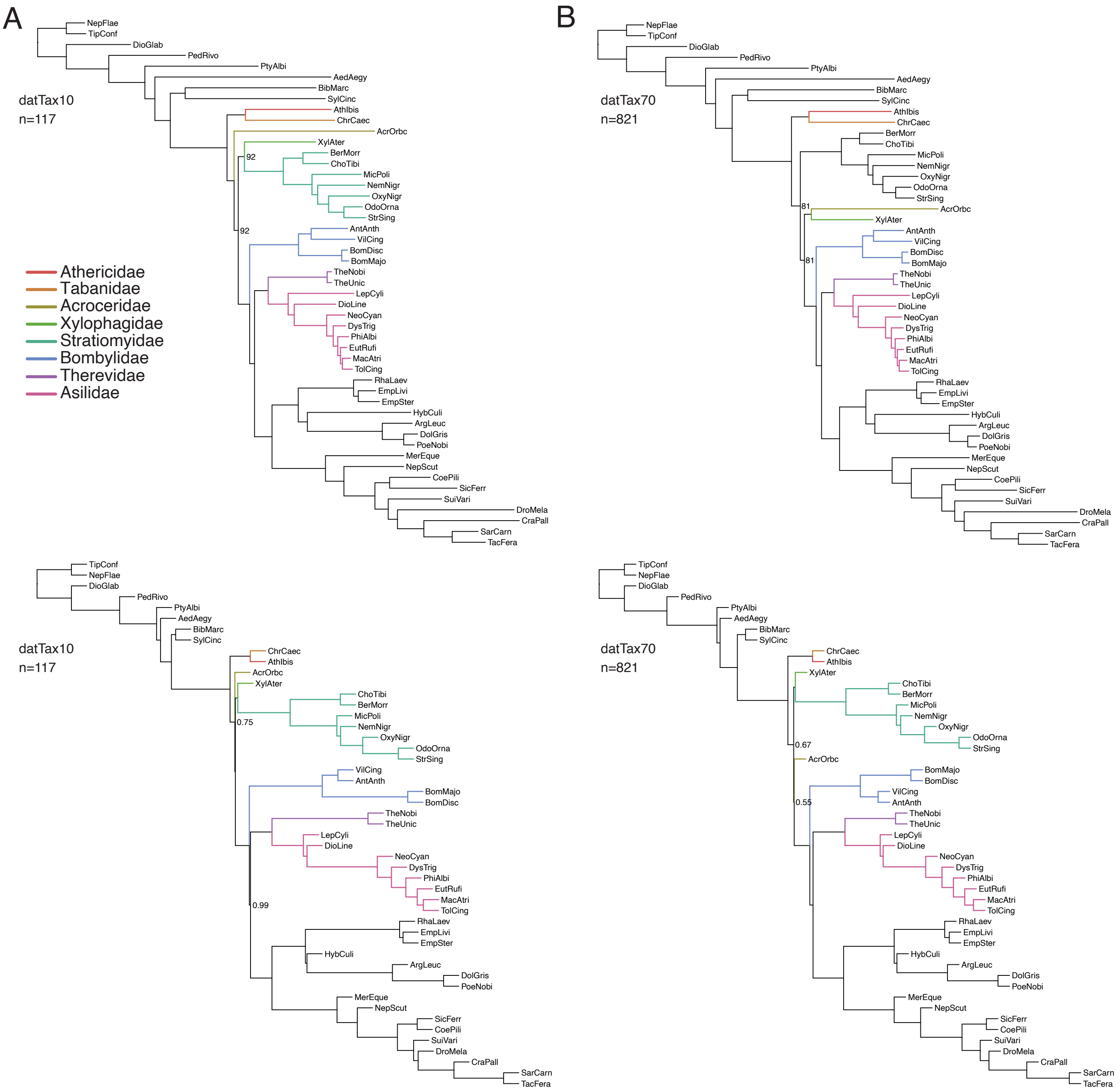


**Supplemental Figure S4: Summary of phylogenetic trees built using the reduced taxon dataset.** **(A)** Phylogenetic trees constructed from the datTax10 dataset using the concatenation based (above) and coalescence based (below) approaches. **(B)** Phylogenetic trees constructed from the datTax70 dataset as described above.

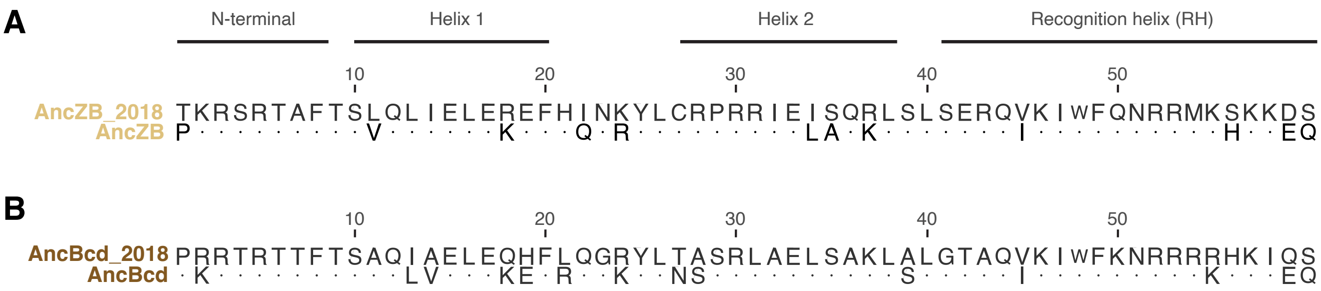


**Supplemental Figure S5: Updated ancestral sequence reconstruction of AncZB and AncBcd.** Comparison of previous AncZB **(A)** and AncBcd **(B)** ancestral sequences obtained from Liu et al. 2018 and those inferred in the current study. Conserved amino acids between alingmnets are given as dots.

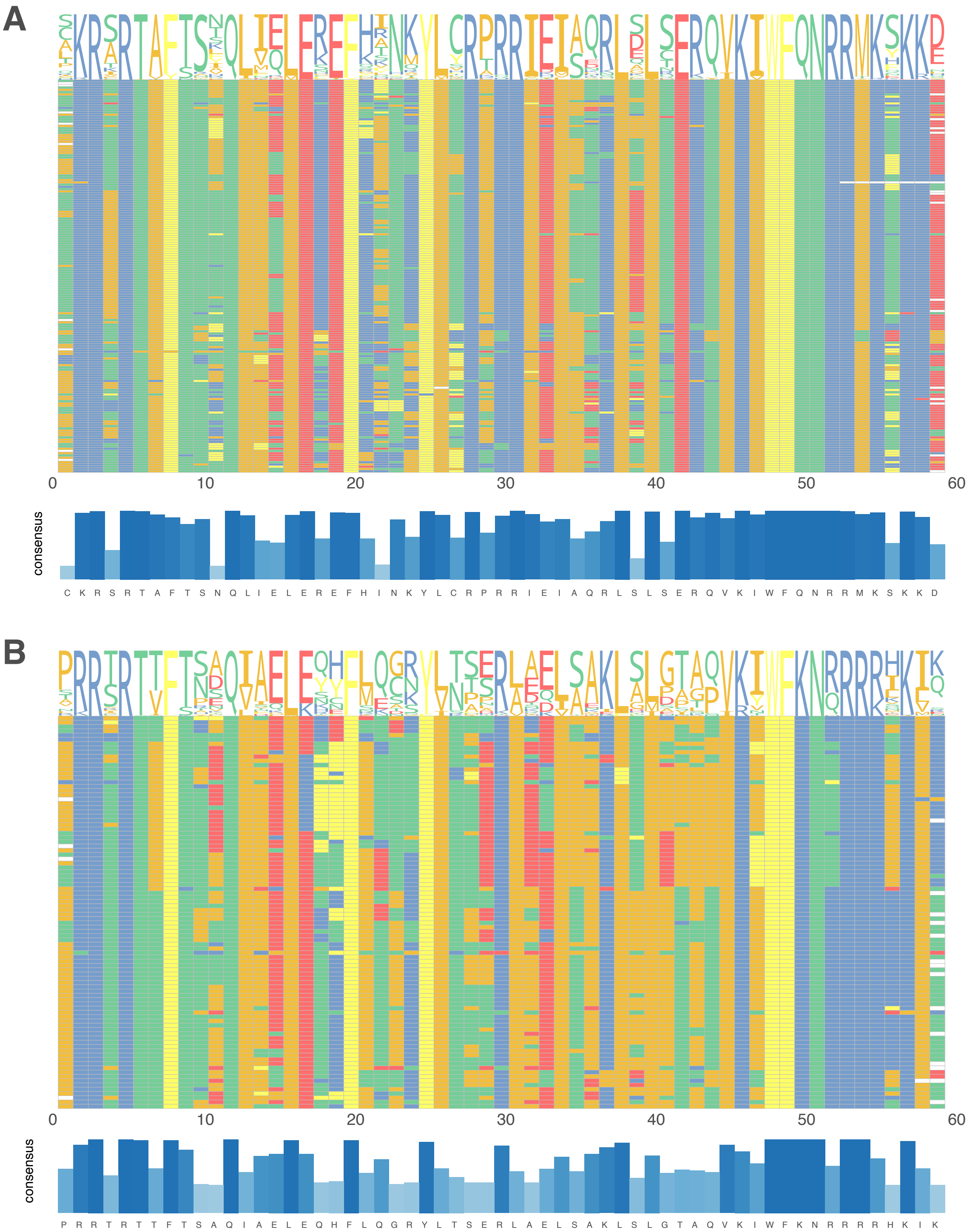


**Supplemental Figure S6: Alignment of Zen and Bcd homeodomains across all species.** Complete alignments of Zen **(A)** and Bcd **(B)** homeodomain sequences from all species in the current study. Amino acids in the alignment are coloured according to their side-chain chemistry. Representative sequence logos for each site are given above the alignment and the consensus conserved percentage of each site is given below as a bar chart.

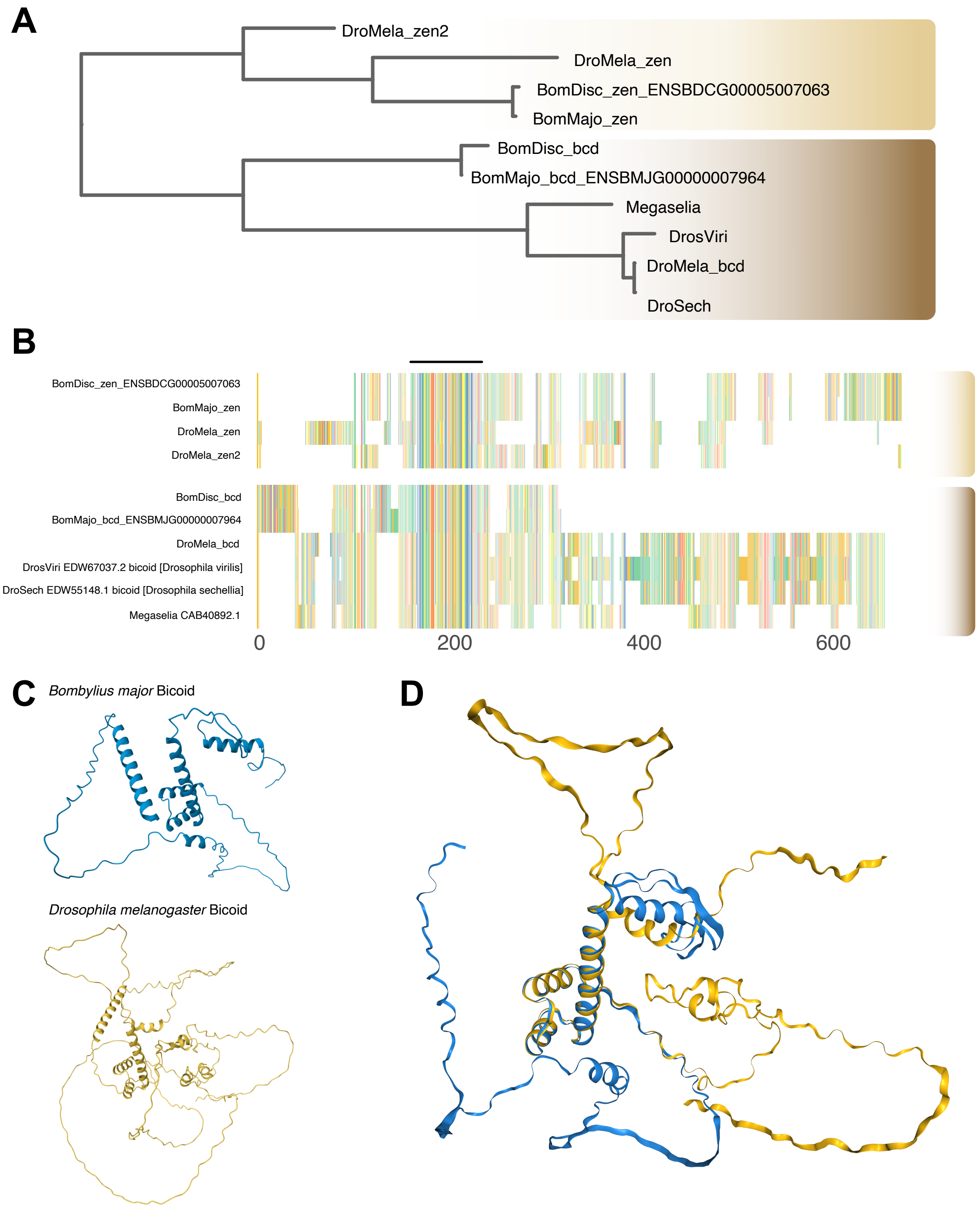


**Supplemental Figure S7: Evolution of complete *bicoid* ORF sequences.** **(A)** Gene tree constructed using alignments of full ORFs of *zen* and *bcd* from *Drosophila melanogaster*, *Drosophila virilis*, *Drosophila sechellia*, *Megaselia abdita*, *Bombylius major*, and *Bombylius discolor*. **(B)** Visualisation of the multiple sequence alignment of *zen* and *bcd* ORFs used in **(A)**. The *zen* sequences are labelled with a yellow bar and *bcd* sequences with a brown bar. Gaps in the alignment are given as white space and the location of the conserved homeodomain is shown with a black bar above the alignment. **(C)** Protein models for *Bombylius major* (blue) and *Drosophila melanogaster* (yellow) *bcd* proteins as inferred by Alphafold. **(D)** Alignment of *Bombylius major* (blue) and *Drosophila melanogaster* (yellow) *bcd* models.

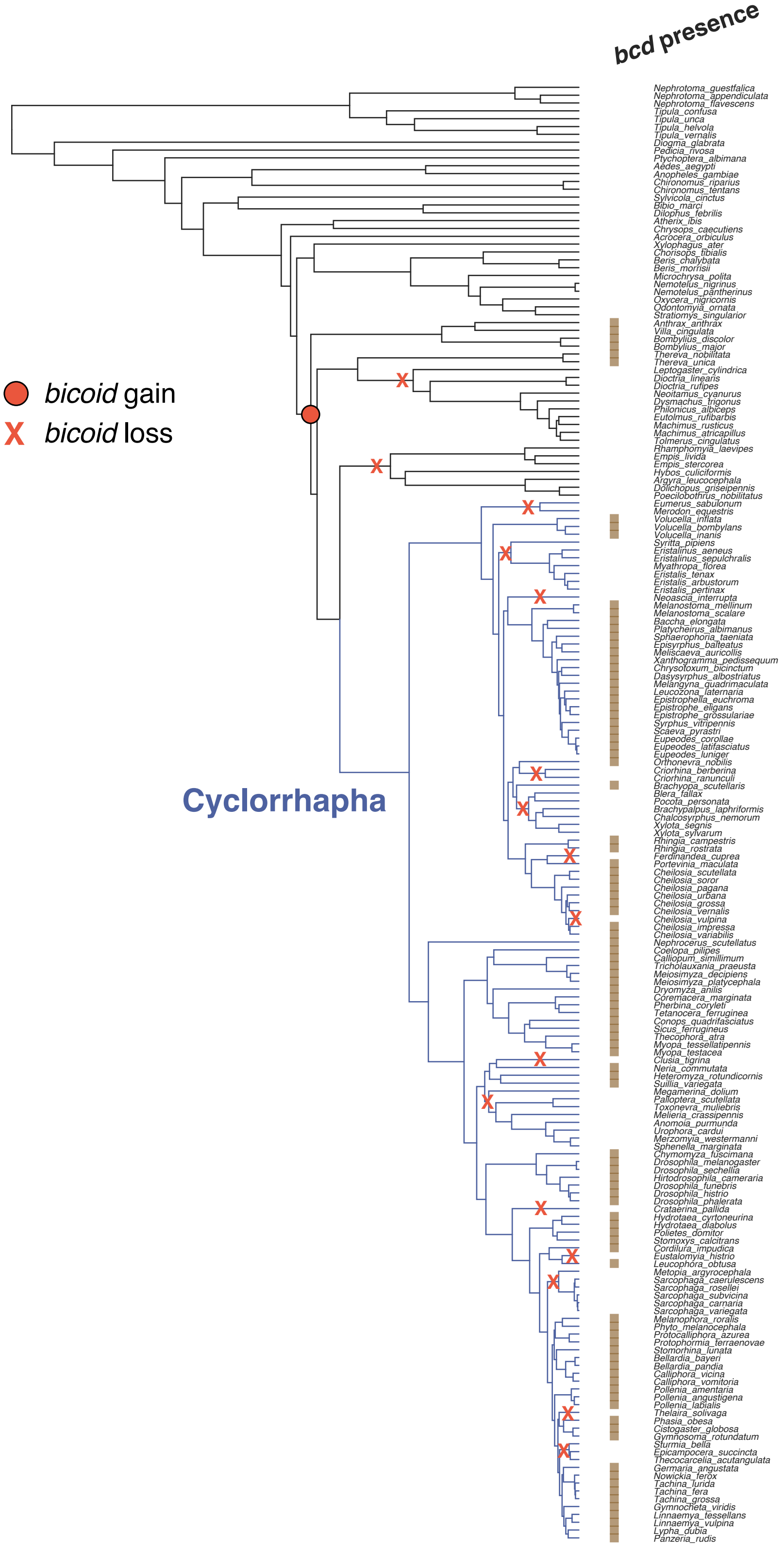


**Supplemental Figure S8: Patterns of *bicoid* loss across flies.** Species tree with *bicoid* presence annotated with brown boxes. Inferred *bicoid* gain (red circle) and loss (red X) events are plotted on tree branches. Cyclorrhapha clade is labelled and colour with blue branches.

### Supplementary Tables

**Table S1: Species sampled and genome source information.**

| Species name | Assembly name | GenBank accession | Reference |
|---|---|---|---|
| *Acrocera orbiculus* | idAcrOrbc1.1 | GCA_947359355.1 | [1] |
| *Aedes aegypti* | AaegL5.0 | GCA_002204515.2 | [2] |
| *Anomoia purmunda* | idAnoPurm1.1 | GCA_951828415.1 | [3] |
| *Anopheles gambiae* | idAnoGambNW-F1-1 | GCA_943734735.2 | [4] |
| *Anthrax anthrax* | idAntAnth1.1 | GCA_963971155.1 | NA |
| *Argyra leucocephala* | idArgLeuc1.1 | GCA_963942445.1 | [5] |
| *Atherix ibis* | idAthIbis2.2 | GCA_958298945.2 | [6] |
| *Baccha elongata* | idBacElon1.2 | GCA_951217065.2 | NA |
| *Bellardia bayeri* | idBelBaye1.1 | GCA_950370525.1 | [7] |
| *Bellardia pandia* | idBelPand1.2 | GCA_916048285.2 | [8] |
| *Beris chalybata* | idBerChal2.1 | GCA_949128065.1 | [9] |
| *Beris morrisii* | idBerMorr1.1 | GCA_951812415.1 | [10] |
| *Bibio marci* | idBibMarc1.2 | GCA_910594885.2 | [11] |
| *Blera fallax* | idBleFall4.1 | GCA_946965025.1 | [12] |
| *Bombylius discolor* | idBomDisc1.1 | GCA_939192795.1 | [13] |
| *Bombylius major* | idBomMajo1.1 | GCA_932526495.1 | [14] |
| *Brachyopa scutellaris* | idBraScut1.1 | GCA_949775065.1 | [15] |
| *Brachypalpus laphriformis* | idBraLaph1.1 | GCA_945910035.1 | [16] |
| *Calliopum simillimum* | idCalSimi1.1 | GCA_951812925.1 | [17] |
| *Calliphora vicina* | idCalVici1.1 | GCA_958450345.1 | [18] |
| *Calliphora vomitoria* | idCalVomi1.2 | GCA_942486065.2 | [19] |
| *Chalcosyrphus nemorum* | idChaNemo1.1 | GCA_949716465.1 | [20] |
| *Cheilosia grossa* | idCheGros1.1 | GCA_963082955.1 | [21] |
| *Cheilosia impressa* | idCheImpr3.1 | GCA_948293265.1 | [22] |
| *Cheilosia pagana* | idChePaga1.1 | GCA_936431705.1 | [23] |
| *Cheilosia scutellata* | idCheScut6.1 | GCA_955612985.1 | [24] |
| *Cheilosia soror* | idCheSoro4.1 | GCA_949372485.1 | [25] |
| *Cheilosia urbana* | idCheUrba1.1 | GCA_946477585.1 | [26] |
| *Cheilosia variabilis* | idCheVari2.1 | GCA_951230905.1 | [27] |
| *Cheilosia vernalis* | idCheVern2.1 | GCA_949126925.1 | [28] |
| *Cheilosia vulpina* | idCheVulp2.1 | GCA_916610125.1 | [29] |
| *Chironomus riparius* | PGI-CHIRRI-v4 | GCA_917627325.4 | NA |
| *Chironomus tentans* | idChiTent1.1 | GCA_963573255.1 | [30] |
| *Chorisops tibialis* | idChoTibi2.1 | GCA_963669355.1 | [31] |
| *Chrysops caecutiens* | idChrCaec1.1 | GCA_963971475.1 | [32] |
| *Chrysotoxum bicinctum* | idChrBici1.1 | GCA_911387755.1 | [33] |
| *Chymomyza fuscimana* | idChyFusc2.1 | GCA_949987675.1 | [34] |
| *Cistogaster globosa* | idCisGlob1.1 | GCA_937654795.1 | [35] |
| *Clusia tigrina* | idCluTigr1.2 | GCA_920105625.2 | [36] |
| *Coelopa pilipes* | idCoePili4.1 | GCA_947389925.1 | [37] |
| *Conops quadrifasciatus* | idConQuad1.1 | GCA_949752815.1 | [38] |

| | | | |
|---|---|---|---|
| *Cordilura impudica* | idCorImpu1.1 | GCA_963682025.1 | NA |
| *Coremacera marginata* | idCorMarg1.1 | GCA_914767935.1 | [39] |
| *Crataerina pallida* | idCraPall2.1 | GCA_949710015.1 | [40] |
| *Criorhina berberina* | idCriBerb1.2 | GCA_917880715.2 | [41] |
| *Criorhina ranunculi* | idCriRanu1.1 | GCA_951813785.1 | [42] |
| *Dasysyrphus albostriatus* | idDasAlbo1.1 | GCA_946251815.1 | [43] |
| *Dilophus febrilis* | idDilFebr1.1 | GCA_958336335.1 | [44] |
| *Dioctria linearis* | idDioLine1.1 | GCA_963930735.1 | [45] |
| *Dioctria rufipes* | idDioRufi1.1 | GCA_963924295.1 | [46] |
| *Diogma glabrata* | idDioGlab1.1 | GCA_963693315.1 | [47] |
| *Dolichopus griseipennis* | idDolGris1.1 | GCA_963082915.1 | [48] |
| *Drosophila funebris* | idDroFune2.1 | GCA_958295475.1 | [49] |
| *Drosophila histrio* | idDroHist2.2 | GCA_958299025.2 | [50] |
| *Drosophila melanogaster* | Release 6 plus ISO1 MT | GCA_000001215.4 | [51] |
| *Drosophila phalerata* | idDroPhal2.2 | GCA_951394115.2 | [52] |
| *Drosophila sechellia* | ASM438219v2 | GCA_004382195.2 | [53] |
| *Dryomyza anilis* | idDryAnil2.1 | GCA_951804985.1 | [54] |
| *Dysmachus trigonus* | idDysTrig1.1 | GCA_949715965.1 | [55] |
| *Empis livida* | idEmpLivi1.1 | GCA_963932195.1 | [56] |
| *Empis stercorea* | idEmpSter1.1 | GCA_949752835.1 | [57] |
| *Epicampocera succincta* | idEpiSucc1.1 | GCA_932526305.1 | [58] |
| *Epistrophe eligans* | idEpiElig2.1 | GCA_951394125.1 | [59] |
| *Epistrophe grossulariae* | idEpiGros1.1 | GCA_929447395.1 | [60] |
| *Epistrophella euchroma* | idEpiEuco1.1 | GCA_947049315.1 | [61] |
| *Episyrphus balteatus* | idEpiBalt1.1 | GCA_945859705.1 | [62] |
| *Eristalinus aeneus* | idEriAene1.1 | GCA_955652365.1 | [63] |
| *Eristalinus sepulchralis* | idEriSepu1.1 | GCA_944738645.1 | [64] |
| *Eristalis arbustorum* | idEriArbu1.1 | GCA_916610145.1 | [65] |
| *Eristalis pertinax* | idEriPert2.1 | GCA_907269125.1 | [66] |
| *Eristalis tenax* | idEriTena2.2 | GCA_905231855.2 | [67] |
| *Eumerus sabulonum* | idEumSabu1.1 | GCA_951905685.1 | [68] |
| *Eupeodes corollae* | idEupCoro1.1 | GCA_945859685.1 | [69] |
| *Eupeodes latifasciatus* | idEupLati1.1 | GCA_920104205.1 | [70] |
| *Eupeodes luniger* | idEupLuni2.1 | GCA_951509635.1 | [71] |
| *Eustalomyia histrio* | idEusHist1.1 | GCA_949748255.1 | [72] |
| *Eutolmus rufibarbis* | idEutRufi1.1 | GCA_963920795.1 | [73] |
| *Ferdinandea cuprea* | idFerCupr2.1 | GCA_963576555.1 | [74] |
| *Germaria angustata* | idGerAngu1.1 | GCA_963681545.1 | [75] |
| *Gymnocheta viridis* | idGymViri1.1 | GCA_956483585.1 | [76] |
| *Gymnosoma rotundatum* | idGymRotn1.2 | GCA_916610165.2 | [77] |
| *Heteromyza rotundicornis* | idHetRotu1.1 | GCA_951394025.1 | [78] |
| *Hirtodrosophila cameraria* | idHirCame2.1 | GCA_949708635.1 | [79] |
| *Hybos culiciformis* | idHybCuli1.1 | GCA_964007475.1 | NA |
| *Hydrotaea cyrtoneurina* | idHydCyrt1.1 | GCA_958296145.1 | [80] |
| *Hydrotaea diabolus* | idHydDiab1.1 | GCA_963513945.1 | [81] |
| *Leptogaster cylindrica* | idLepCyli1.1 | GCA_963082835.1 | [82] |
| *Leucophora obtusa* | idLeuObtu2.1 | GCA_949987735.1 | [83] |

| | | | |
|---|---|---|---|
| *Leucozona laternaria* | idLeuLate1.2 | GCA_932273885.2 | [84] |
| *Linnaemya tessellans* | idLinTess1.1 | GCA_951800035.1 | [85] |
| *Linnaemya vulpina* | idLinVulp1.1 | GCA_963675445.1 | [86] |
| *Lypha dubia* | idLypDubi1.1 | GCA_947311025.1 | [87] |
| *Machimus atricapillus* | idMacAtri3.1 | GCA_933228815.1 | [88] |
| *Machimus rusticus* | idMacRust1.1 | GCA_951509405.1 | [89] |
| *Megamerina dolium* | idMegDoli1.1 | GCA_963854835.1 | [90] |
| *Meiosimyza decipiens* | idLycDeci1.1 | GCA_963680825.1 | [91] |
| *Meiosimyza platycephala* | idLycPlat1.1 | GCA_963662105.1 | NA |
| *Melangyna quadrimaculata* | idMelQuad1.1 | GCA_949320155.1 | [92] |
| *Melanophora roralis* | idMelRora1.1 | GCA_963583895.1 | [93] |
| *Melanostoma mellinum* | idMelMell2.1 | GCA_914767635.1 | [94] |
| *Melanostoma scalare* | idMelScal2.1 | GCA_949752695.1 | [95] |
| *Melieria crassipennis* | idMelCras1.1 | GCA_963668005.1 | [96] |
| *Meliscaeva auricollis* | idMelAuri2.1 | GCA_948107695.1 | [97] |
| *Merodon equestris* | idMerEque2.1 | GCA_958301585.1 | [98] |
| *Merzomyia westermanni* | idMerWest1.1 | GCA_949987695.1 | [99] |
| *Metopia argyrocephala* | idMetArgy1.1 | GCA_963576795.1 | [100] |
| *Microchrysa polita* | idMicPoli1.1 | GCA_949715475.1 | [101] |
| *Myathropa florea* | idMyaFlor2.1 | GCA_930367185.1 | [102] |
| *Myopa tessellatipennis* | idMyoTess1.2 | GCA_943737955.2 | [103] |
| *Myopa testacea* | idMyoTest1.1 | GCA_949629155.1 | [104] |
| *Nemotelus nigrinus* | idNemNigr1.1 | GCA_947369275.1 | [105] |
| *Nemotelus pantherinus* | idNemPant2.1 | GCA_963978885.1 | [106] |
| *Neoascia interrupta* | idNeoInte1.1 | GCA_947623515.1 | [107] |
| *Neoitamus cyanurus* | idNeoCyan1.1 | GCA_947538895.1 | [108] |
| *Nephrocerus scutellatus* | idNepScut1.1 | GCA_947095585.1 | [109] |
| *Nephrotoma appendiculata* | idNepAppe1.1 | GCA_947310385.1 | [110] |
| *Nephrotoma flavescens* | idNepFlae1.1 | GCA_932526605.1 | [111] |
| *Nephrotoma guestfalica* | idNepGues1.1 | GCA_963691935.1 | [112] |
| *Neria commutata* | idNerComm1.1 | GCA_963457695.1 | [113] |
| *Nowickia ferox* | idNowFero1.1 | GCA_936439885.1 | [114] |
| *Odontomyia ornata* | idOdoOrna1.1 | GCA_963969285.1 | [115] |
| *Orthonevra nobilis* | idOrtNobi1.1 | GCA_963555765.1 | [116] |
| *Oxycera nigricornis* | idOxyNigr1.1 | GCA_963942545.1 | [117] |
| *Palloptera scutellata* | idPalScut3.1 | GCA_958295655.1 | [118] |
| *Panzeria rudis* | idPanRudi1.1 | GCA_956483635.1 | [119] |
| *Pedicia rivosa* | idPedRivo1.1 | GCA_963082725.1 | NA |
| *Phasia obesa* | idPhaObes1.1 | GCA_949628195.1 | [120] |
| *Pherbina coryleti* | idPheCory1.1 | GCA_943735915.1 | [121] |
| *Philonicus albiceps* | idPhiAlbi1.1 | GCA_963969385.1 | [122] |
| *Phyto melanocephala* | idPhyMeln1.1 | GCA_941918925.1 | [123] |
| *Platycheirus albimanus* | idPlaAlba1.2 | GCA_916050605.2 | [124] |
| *Pocota personata* | idPocPers1.1 | GCA_963082735.1 | [125] |
| *Poecilobothrus nobilitatus* | idPoeNobi2.1 | GCA_947095535.1 | [126] |
| *Polietes domitor* | idPolDomi1.1 | GCA_947397865.1 | [127] |
| *Pollenia amentaria* | idPolAmen1.1 | GCA_943735925.1 | [128] |

| | | | |
|---|---|---|---|
| *Pollenia angustigena* | idPolAngu1.1 | GCA_930367215.1 | [129] |
| *Pollenia labialis* | idPolLabi1.1 | GCA_949318255.1 | [130] |
| *Portevinia maculata* | idPorMacu1.1 | GCA_949715645.1 | [131] |
| *Protocalliphora azurea* | idProAzur1.1 | GCA_932274085.1 | [132] |
| *Protophormia terraenovae* | idProTerr1.1 | GCA_951394005.1 | [133] |
| *Ptychoptera albimana* | idPtyAlbi3.1 | GCA_961205885.1 | [134] |
| *Rhamphomyia laevipes* | idRhaLaev1.1 | GCA_963920725.1 | [135] |
| *Rhingia campestris* | idRhiCamp1.1 | GCA_932526625.1 | [136] |
| *Rhingia rostrata* | idRhiRost1.1 | GCA_949824845.1 | [137] |
| *Sarcophaga caerulescens* | idSarCaer1.1 | GCA_927399465.1 | [138] |
| *Sarcophaga carnaria* | idSarCarn1.1 | GCA_958299815.1 | [139] |
| *Sarcophaga rosellei* | idSarRose1.1 | GCA_930367235.1 | [140] |
| *Sarcophaga subvicina* | idSarSubv1.2 | GCA_936449025.2 | [141] |
| *Sarcophaga variegata* | idSarVari1.1 | GCA_932273835.1 | [142] |
| *Scaeva pyrastri* | idScaPyra1.1 | GCA_905146935.1 | [143] |
| *Sicus ferrugineus* | idSicFerr1.1 | GCA_922984085.1 | [144] |
| *Sphaerophoria taeniata* | idSphTaen1.1 | GCA_943590905.1 | [145] |
| *Sphenella marginata* | idSphMarg1.1 | GCA_951509765.1 | [146] |
| *Stomorhina lunata* | idStoLuna1.1 | GCA_933228675.1 | [147] |
| *Stomoxys calcitrans* | idStoCalc2.1 | GCA_963082655.1 | [148] |
| *Stratiomys singularior* | idStrSing1.1 | GCA_954870665.1 | [149] |
| *Sturmia bella* | idStuBell1.1 | GCA_963662145.1 | [150] |
| *Suillia variegata* | idSuiVari3.1 | GCA_949127995.1 | [151] |
| *Sylvicola cinctus* | idSylCinc2.1 | GCA_963854165.1 | [152] |
| *Syritta pipiens* | idSyrPipi1.1 | GCA_905187475.1 | [153] |
| *Syrphus vitripennis* | idSyrVitr1.1 | GCA_958431115.1 | [154] |
| *Tachina fera* | idTacFera2.1 | GCA_905220375.1 | [155] |
| *Tachina grossa* | idTacGros1.1 | GCA_949987645.1 | [156] |
| *Tachina lurida* | idTacLuri1.1 | GCA_944452675.1 | [157] |
| *Tetanocera ferruginea* | idTetFerr1.1 | GCA_958299015.1 | [158] |
| *Thecocarcelia acutangulata* | idTheAcut1.1 | GCA_914767995.1 | [159] |
| *Thecophora atra* | idTheAtra2.1 | GCA_937620795.1 | [160] |
| *Thelaira solivaga* | idTheSoli1.1 | GCA_947397855.1 | [161] |
| *Thereva nobilitata* | idTheNobi1.1 | GCA_963855945.1 | [162] |
| *Thereva unica* | idTheUnic2.1 | GCA_949987705.1 | [163] |
| *Tipula confusa* | idTipConf1.1 | GCA_963556175.1 | [164] |
| *Tipula helvola* | idTipHelv1.1 | GCA_963556165.1 | [165] |
| *Tipula unca* | idTipUnca1.1 | GCA_951394425.1 | [166] |
| *Tipula vernalis* | idTipVern1.1 | GCA_958295665.1 | [167] |
| *Tolmerus cingulatus* | idTolCing1.1 | GCA_959613345.1 | [168] |
| *Toxonevra muliebris* | idToxMuli2.1 | GCA_963691655.1 | [169] |
| *Tricholauxania praeusta* | idTriPrae1.1 | GCA_949775025.1 | [170] |
| *Urophora cardui* | idUroCard1.1 | GCA_960531455.1 | [171] |
| *Villa cingulata* | idVilCing2.1 | GCA_951394055.1 | [172] |
| *Volucella bombylans* | idVolBomb1.1 | GCA_949129095.1 | [173] |
| *Volucella inanis* | idVolInan1.1 | GCA_907269105.1 | [174] |
| *Volucella inflata* | idVolInfl1.1 | GCA_928272305.1 | [175] |

| | | | |
|---|---|---|---|
| *Xanthogramma pedissequum* | idXanPedi1.1 | GCA_910595825.1 | [176] |
| *Xylophagus ater* | idXylAter1.1 | GCA_963422695.1 | NA |
| *Xylota segnis* | idXylSegn2.1 | GCA_963583995.1 | [177] |
| *Xylota sylvarum* | idXylSylv2.1 | GCA_905220385.1 | [178] |

11. Sivell O, Sivell D, Natural History Museum Genome Acquisition Lab, Darwin Tree of Life Barcoding collective, Wellcome Sanger Institute Tree of Life programme, Wellcome Sanger Institute Scientific Opera-

tions: DNA Pipelines collective, et al. The genome sequence of the St Mark's fly, Bibio marci (Linnaeus, 1758). Wellcome Open Res. 2021;6: 285.

23. Falk S, Poole O, University of Oxford and Wytham Woods Genome Acquisition Lab, Darwin Tree

of Life Barcoding collective, Wellcome Sanger Institute Tree of Life Management, Samples and Laboratory team, Wellcome Sanger Institute Scientific Operations: Sequencing Operations, et al. The genome sequence of the parsley Cheilosia, Cheilosia pagana (Meigen, 1822). Wellcome Open Res. 2024;9: 54.

35. Falk S, Lennon R, University of Oxford Genome and Wytham Woods Genome Acquisition Lab, Darwin Tree of Life Barcoding collective, Wellcome Sanger Institute Tree of Life programme, Wellcome

Sanger Institute Scientific Operations: DNA Pipelines collective, et al. The genome sequence of a tachinid fly, Cistogaster globosa (Fabricius, 1775). Wellcome Open Res. 2023;8: 462.

47. McCulloch J, Crowley LM, University of Oxford and Wytham Woods Genome Acquisition Lab, Darwin Tree of Life Barcoding collective, Wellcome Sanger Institute Tree of Life Management, Samples and Laboratory team, Wellcome Sanger Institute Scientific Operations: Sequencing Operations, et al. The

genome sequence of a cranefly, Diogma glabrata (Meigen, 1818). Wellcome Open Res. 2025;10: 20.

63. Sivell O, Raper C, Mitchell R, Sivell D, Natural History Museum Genome Acquisition Lab, Darwin

Tree of Life Barcoding collective, et al. The genome sequence of a hoverfly Eristalinus aeneus (Scopoli, 1763). Wellcome Open Res. 2024;9: 69.

94. Hawkes W, Wotton K, University of Oxford and Wytham Woods Genome Acquisition Lab, Darwin Tree of Life Barcoding collective, Wellcome Sanger Institute Tree of Life programme, Wellcome Sanger Institute Scientific Operations: DNA Pipelines collective, et al. The genome sequence of the dumpy grass

hoverfly, Melanostoma mellinum (Linnaeus, 1758). Wellcome Open Res. 2022;7: 59.

108. Crowley LM, Akinmusola RY, University of Oxford and Wytham Woods Genome Acquisition Lab, Darwin Tree of Life Barcoding collective, Wellcome Sanger Institute Tree of Life Management, Samples

and Laboratory team, Wellcome Sanger Institute Scientific Operations: Sequencing Operations, et al. The genome sequence of the common awl robberfly, Neoitamus cyanurus (Loew, 1849). Wellcome Open Res. 2024;9: 289.

124. Crowley LM, Woodcock KJ, University of Oxford and Wytham Woods Genome Acquisition Lab,

Darwin Tree of Life Barcoding collective, Wellcome Sanger Institute Tree of Life Management, Samples and Laboratory team, Wellcome Sanger Institute Scientific Operations: Sequencing Operations, et al. The genome sequence of the white-footed hoverfly, Platycheirus albimanus (Fabricius, 1781). Wellcome Open Res. 2023;8: 572.

140. Falk S, University of Oxford and Wytham Woods Genome Acquisition Lab, Darwin Tree of Life Barcoding collective, Wellcome Sanger Institute Tree of Life programme, Wellcome Sanger Institute Scientific Operations: DNA Pipelines collective, Tree of Life Core Informatics collective, et al. The genome of Roselle's

flesh fly Sarcophaga ( Helicophagella) rosellei (Böttcher, 1912). Wellcome Open Res. 2023;8: 43.
